# Loss of *MEF2C* function by enhancer mutation leads to neuronal mitochondria dysfunction and motor deficits in mice

**DOI:** 10.1101/2024.07.15.603186

**Authors:** Ali Yousefian-Jazi, Suhyun Kim, Seung-Hye Choi, Jiyeon Chu, Phuong Thi-Thanh Nguyen, Uiyeol Park, Kayeong Lim, Hongik Hwang, Kyungeun Lee, Yeyun Kim, Seung Jae Hyeon, Hyewhon Rhim, Hannah L. Ryu, Grewo Lim, Thor D. Stein, Hoon Ryu, Junghee Lee

## Abstract

Genetic changes and epigenetic modifications are associated with neuronal dysfunction in the pathogenesis of neurodegenerative disorders. However, the mechanism behind genetic mutations in the non-coding region of genes that affect epigenetic modifications remains unclear. Here, we identified an ALS-associated SNP located in the intronic region of *MEF2C* (rs304152), residing in a putative enhancer element, using convolutional neural network. The enhancer mutation of *MEF2C* reduces own gene expression and consequently impairs mitochondrial function in motor neurons. MEF2C localizes and binds to the mitochondria DNA, and directly modulates mitochondria-encoded gene expression. CRISPR/Cas-9-induced mutation of the *MEF2C* enhancer decreases expression of mitochondria-encoded genes. Moreover, *MEF2C* mutant cells show reduction of mitochondrial membrane potential, ATP level but elevation of oxidative stress. *MEF2C* deficiency in the upper and lower motor neurons of mice impairs mitochondria-encoded genes, and leads to mitochondrial metabolic disruption and progressive motor behavioral deficits. Together, *MEF2C* dysregulation by the enhancer mutation leads to mitochondrial dysfunction and oxidative stress, which are prevalent features in motor neuronal damage and ALS pathogenesis. This genetic and epigenetic crosstalk mechanism provides insights for advancing our understanding of motor neuron disease and developing effective treatments.

## Introduction

Amyotrophic lateral sclerosis (ALS), also known as Lou Gehrig’s disease, is a progressive neurodegenerative disease characterized by the loss of motor neurons in both the upper and lower spinal cord. The loss of motor neurons leads to muscle wasting and paralysis. While around 90-95% of ALS cases are sporadic, the remaining 5-10% of cases are familial. Among familial ALS cases with a family history, roughly 20% are caused by mutations in the *SOD1* gene, the first one discovered to be involved in ALS. Additional genes linked to familial ALS including *TARDBP* and *FUS*, accounting for 4-5% of cases each. Notably, mutations in *C9ORF72* contribute to nearly 40% of familial cases, while mutations in other unidentified genes cause the remainder (1, 2). Studies suggest that epigenetic factors play a role in the etiology of ALS, as well as other neurodegenerative and neuropsychiatric diseases (3–8). Epigenetic modifications can influence various cellular processes, including mitochondrial function, energy metabolism, oxidative stress, and programmed cell death, by regulating gene expression (9). However, the precise mechanisms by which genetic mutations in the non-coding regions of genes can trigger epigenetic modifications and contribute to the neuropathology of ALS remain incompletely understood.

Genome-wide association study (GWAS) has become a solid method for identifying intronic and intergenic genetic variations linked to various complex diseases. For ALS, GWAS has successfully pointed to genes like *C9ORF72*, *SARM*, *UNC13A, SCFD1*, *TBK1*, *MOBP*, *C21ORF2*, and *KIF5A* as potential risk factors (10–12). However, even though GWAS has revealed many promising leads, it can be challenging to pinpoint the exact single-nucleotide polymorphisms (SNPs) responsible for the disease, especially when they are rare and their effects are subtle (13). This limitation has spurred the development of post-GWAS analysis method, such as those using convolutional neural networks (CNNs) trained on epigenetic data (14). These new approaches aim to identify functional, but rare, risk-variants hidden within non-coding regions of the genome. One such improved CNN model constructed with uncertain class labels has been applied to the GWAS meta-analysis from 14,791 ALS cases and 26,898 healthy controls. This analysis successfully identified two potential risk-SNPs on chromosomes 3 and 17, opening the door for further investigation (10, 15).

Building on the largest ALS GWAS ever conducted, which encompasses data from over 20,000 patients and 59,000 healthy individuals, we scrutinized deeper into understanding the genetic roots of this devastating disease. We further examined genetic correlations between ALS and schizophrenia (SCZ) by analyzing a separate GWAS. This combined approach led us to identify four promising risk-SNPs, all hidden within the non-coding region of a gene called myocyte enhancer factor 2C (*MEF2C*). Intrigued by these findings, we explored whether these SNPs affect MEF2C’s activity. Interestingly, we discovered that MEF2C localized to the mitochondria of motor neurons and modulates mitochondrial DNA transcription and mitochondrial function. Lastly, we verified how a loss of *MEF2C* function contributes to neurodegeneration and motor behavior deficit in mice.

## Results

### Identification and biological characterization of ALS risk-SNPs

To identify the functional risk-SNPs using a CNN model based on the genetic correlation between ALS and SCZ, we used genetic association data from the large-scale GWAS meta-analyses for ALS and SCZ (Fig. 1A). The ALS GWAS data includes 10,031,630 imputed variants from 20,806 patients diagnosed with ALS and 59,804 control subjects (12), and the SCZ GWAS data is identified on 36,989 cases and 113,075 controls (16). We trained the CNN model with uncertain labels on the extracted epigenetic features map of SNPs in association and control blocks (Methods). The model’s performance was shown by the area under the curve on the test blocks for ALS and SCZ equal to 0.92 and 0.96, respectively (Fig. 1B). The CNN model predicts a prediction_score between 0 and 1 for each SNP, and those with prediction_score > 0.5 considered as the initial candidates of risk-SNPs in each block (Fig. 1C). Among 2141 and 7562 SNPs with prediction_score > 0.5 in ALS and SCZ, respectively, 669 and 2466 SNPs had been selected based on the highest prediction score in each block. Then, the target genes were assigned to each SNP using the gene body, the 3*kb* upstream of the transcription start site as a promoter region and LASSO transcriptional enhancers in brain cell lines as an enhancer region (17). We identified 44 genes shared between ALS and SCZ (Fig. 1A). Then, we searched for candidate ALS risk-SNPs that shared the same linkage disequilibrium (LD) with SCZ-associated risk-SNPs linked to 44 genes (18). Finally, we arrived at the list of 27 SNPs and 12 linked gene candidates associated with ALS (Supplementary Table 1). Among the 12 genes, *MEF2C* was reduced by more than 50% in frontal cortex mRNA transcriptome data from 22-month-old ALS-FUS male mice(19) (Supplementary Fig. 1). Furthermore, decreased MEF2C immunoreactivity was observed by immunofluorescence staining in cortical pyramidal neurons of ALS patients and ALS mice model (G93A) (Supplementary Fig. 2A, C). Similarly, MEF2C immunoreactivity level significantly decreased in lower motor neurons in the lumbar spinal cord of both ALS patients and the G93A mouse model (Supplementary Fig. 2B, D). Four candidate risk-SNPs, rs700587, rs304151, rs304152 and rs304153, located in the intronic region of the *MEF2C* gene, were identified as the risk-SNPs by our model. The rs304152 located in H3K27ac peak region which is labeled as an active enhancer marker(20), and is in the strong LD with other three SNPs, as well (Fig. 1D-E). The Hi-C data across human prefrontal cortex indicates the potential long-range chromatin interactions between rs304152 and *MEF2C* promoter region (Supplementary Fig. 3) (21). Moreover, Genotype-Tissue Expression (GTEx) database (https://gtexportal.org) illustrates the lower *MEF2C* expression level for the risk-allele G of rs304152 in the most brain regions (Fig. 1F). Therefore, rs304152 and *MEF2C* was chosen as the ALS-associated SNP and gene for the further analysis in this study.

**Figure 1.**
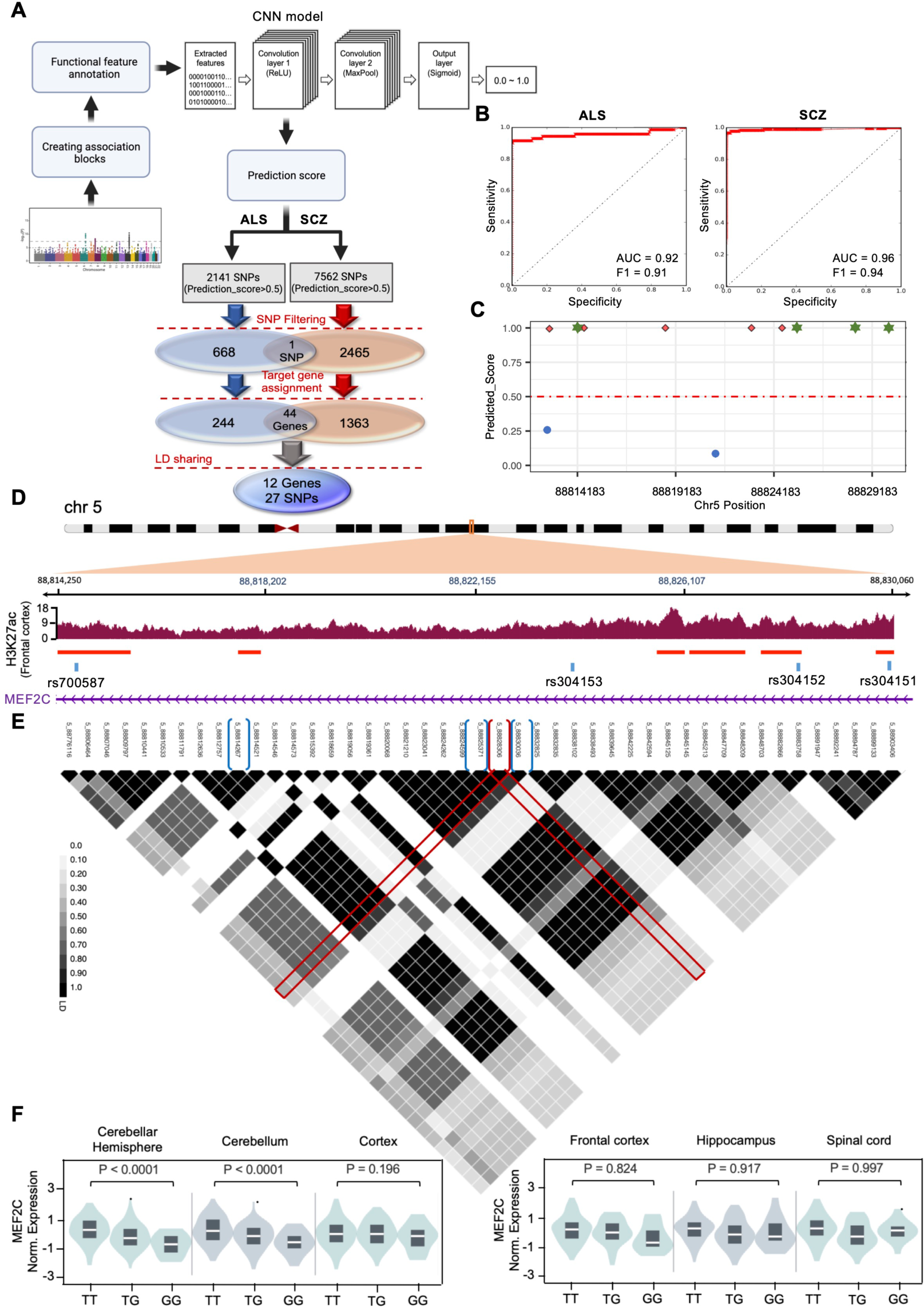
Prioritizing rs304152 as a potential functional SNP in intronic region of *MEF2C* associated with ALS. **(A)** A research flow starting from ALS and SCZ GWAS meta-analysis data and extracting functional features for each SNP, then using CNN model to initialize the candidate risk-SNPs followed by filtering, target gene assignment and LD sharing steps. **(B)** The CNN model performance for prediction of testing blocks for ALS and SCZ model. **(C)** The prediction score for each SNP in the association block contains *MEF2C* gene body. Green stars are candidate SNPs. **(D)** Active regulatory regions coverage and peaks marked by H3K27ac in frontal cortex extracted from GTEX IGV browser. **(E)** The LD plots for the region from 88.7 to 88.9 Mb in chromosome 5 including candidate SNPs. LD block structure was estimated with the Haploview software. The red brocket, rs304152, is the chosen SNP which has the high LD scores with other blue brocket candidate risk-SNPs, rs700587, rs304153 and rs304151. **(F)** Violin plots of the *MEF2C* normalized expression level according to alleles of rs304152 in different brain regions. The information was extracted from GTEX database.

### Mutation in the enhancer of *MEF2C*, rs304152, reduces *MEF2C* mRNA transcription by impairing ATF4 transcription factor binding

Since the mutation site of rs304152 resides in a potential enhancer region, we hypothesized that this mutation may affect the promoter activity of its own gene. In this context, we generated plasmid constructs in which a duplex of a 21-bp intron sequence carrying the major or minor allele residue in the SNP-corresponding position was coupled to the human *MEF2C* promoter and downstream luciferase reporter. Given the fact that the mutation site of the *MEF2C* enhancer harbors an ATF4 binding element, the cells were transfected with ATF4 for 48hrs, and then the *MEF2C* promoter activity was measured using the Dual-Luciferase reporter assay system (Promega) (Fig. 2A). The rs304152-G allele (MT) showed less luciferase activity than the rs304152-T allele (WT) on the *MEF2C* promoter in NSC-34 cells (Fig. 2B, Supplementary Fig. 4A). The cells transfected by ATF4 revealed higher luciferase activity for both WT and MT, indicated the activator role of ATF4 binding to the region residing rs304152. To confirm the ATF4 occupancy in the *MEF2C* enhancer, we performed ChIP assay using the ChIP grade anti-ATF4 monoclonal antibody followed by qPCR with primers targeting the enhancer region (Fig. 2C). ChIP-qPCR results demonstrated the less occupancy of ATF4 in the MT form of *MEF2C* enhancer region (Fig. 2D). Additionally, DNA sequencing of ATF4-ChIP product confirmed that ATF4 preferentially binds to the rs304152-T allele compared to the rs304152-G allele in the MEF2C enhancer (Supplementary Fig. 4B). For further clarification on the activator role of ATF4 for *MEF2C* transcription, enforced overexpression of ATF4 was carried out by transfection of ATF4 plasmid in NSC-34 cells for 12hrs. Using immunofluorescence, we confirmed that the MEF2C immunoreactivity was elevated in the cytosol compartment by increasing ATF4 immunoreactivity level (Fig. 2E). The scatter plot illustrated a high positive correlation (R^2^ = 0.89) between ATF4 and MEF2C immunoreactivity (Fig. 2F). Overall, these findings demonstrate ATF4 transcription factor preferentially bind to the WT form of *MEF2C* enhancer and promotes *MEF2C* transcription (Fig. 2G).

**Figure 2.**
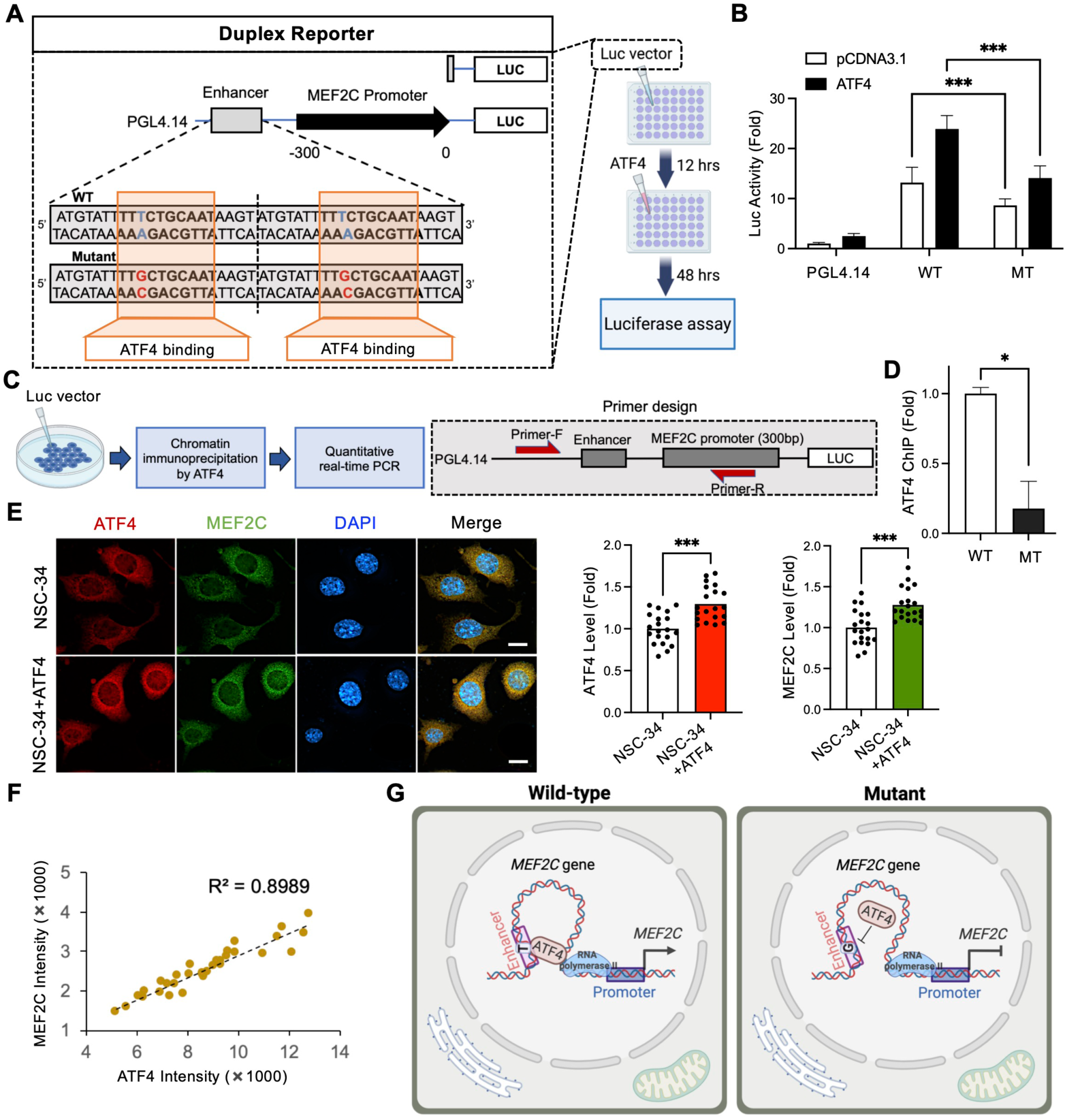
Risk-SNP (rs304152) of *MEF2C* gene significantly reduces the transcription of own gene by inhibiting ATF4 binding to the enhancer region. **(A)** Schematic representation of constructs used in the luciferase reporter assays. **(B)** The construct containing rs304152-G allele (WT) showed approximately 1.7-fold less luciferase activity than the construct containing rs304152-T allele (MT). The experiment was repeated three times. *\*\*\*P* < 0.001; Two-way ANOVA test (*n* = 12/group, *df* = 66). **(C)** A scheme illustrating ChIP-qPCR assay for determining allele-specific DNA–protein interactions of WT and MT form of *MEF2C* enhancer region and anti-ATF4 antibody in NSC-34 cells. **(D)** ChIP-qPCR results showed less ATF4 occupancy at the MT form of *MEF2C* enhancer. *\*P* < 0.05; Student’s t test (*n* = 3/group, *df* = 4). The P values determined by Student’s T-test. **(E)** Immunofluorescence staining for ATF4 (red) and MEF2C (green) in NSC-34 cells transfected by ATF4 for 12 hrs. Right: quantitation of ATF4 and MEF2C immunoreactivity levels. A total of 42 cells count, 7 cells/well, *N* = 3 (NSC-34, NSC-34+ATF4). *\*\*\*P* < 0.001; Student’s t test (*n* = 21/group, *df* = 40). **(F)** The scatter plot showed ATF4 and MEF2C immunoreactivity levels correlation in the cells. **(G)** A scheme summarizing that the mutant form of *MEF2C* enhancer resulted *MEF2C* transcriptional impairment by inhibiting ATF4 transcription factor binding to the enhancer region. Scheme created with BioRender.com. Error bars indicate means ± s.e.m.

### Enhancer mutation, rs304152, impaired *MEF2C* transcription and mitochondrial function

To investigate the functional consequences of rs304152, we generated HEK293T cells carrying the major or minor allele by CRISPR-Cas9 (Fig. 3A). Consistent with our hypothesis, rs304152-G (MT) cells displayed significantly lower *MEF2C* mRNA levels compared to rs304152-T (WT) cells (Fig. 3B). After verifying that the rs304152 mutation decreases *MEF2C* expression level, we investigated which subcellular organelle MEF2C localized in the cytosol to elucidate the specific role of MEF2C in neuron. By observing the immunofluorescence co-staining of MEF2C and TOMM20, an outer mitochondrial membrane marker, we found MEF2C localized to mitochondria in postmortem human cortex pyramidal neurons. The 3D reconstruction and line measurement results confirmed the localization of MEF2C in mitochondria (Fig. 3C). Even though mouse cell line and human tissue show a different pattern of MEF2C localization in the subcellular compartments, we found that a higher expression level of MEF2C is apparently found in the mitochondrial compartment than in the nuclear compartment (Fig. 2E and Fig. 3C). According to the quantification data, 62.7% of TOMM20- and MEF2C-double positive signals are localized to the mitochondria of cortical layer V pyramidal neurons (Supplementary Fig. 5A). We further quantitated MEF2C signals in DAB-stained cells and confirmed that approximately twice as much of MEF2C immunoreactivity is found in the cytosol compartment compared to the nuclear compartment of the cortical layer V pyramidal neurons in both normal and ALS patients (Supplementary Fig. 5B). MEF2C localization in mitochondria was observed in mouse cortical pyramidal neurons as well (Supplementary Fig. 5C). Moreover, we obtained subcellular fractions from mouse brain tissue by sucrose density centrifugation (Fig. 3D). Western blot analysis confirmed that MEF2C exist in enriched mitochondrial fractions of mouse brain tissue (Fig. 3E, Supplementary Fig. 5D). The results showed the LaminB1 and COX4 immunoreactivity mostly in nuclear and mitochondrial fractions, respectively, which indicates the mitochondrial fractions without contamination with other subcellular organelles.

**Figure 3.**
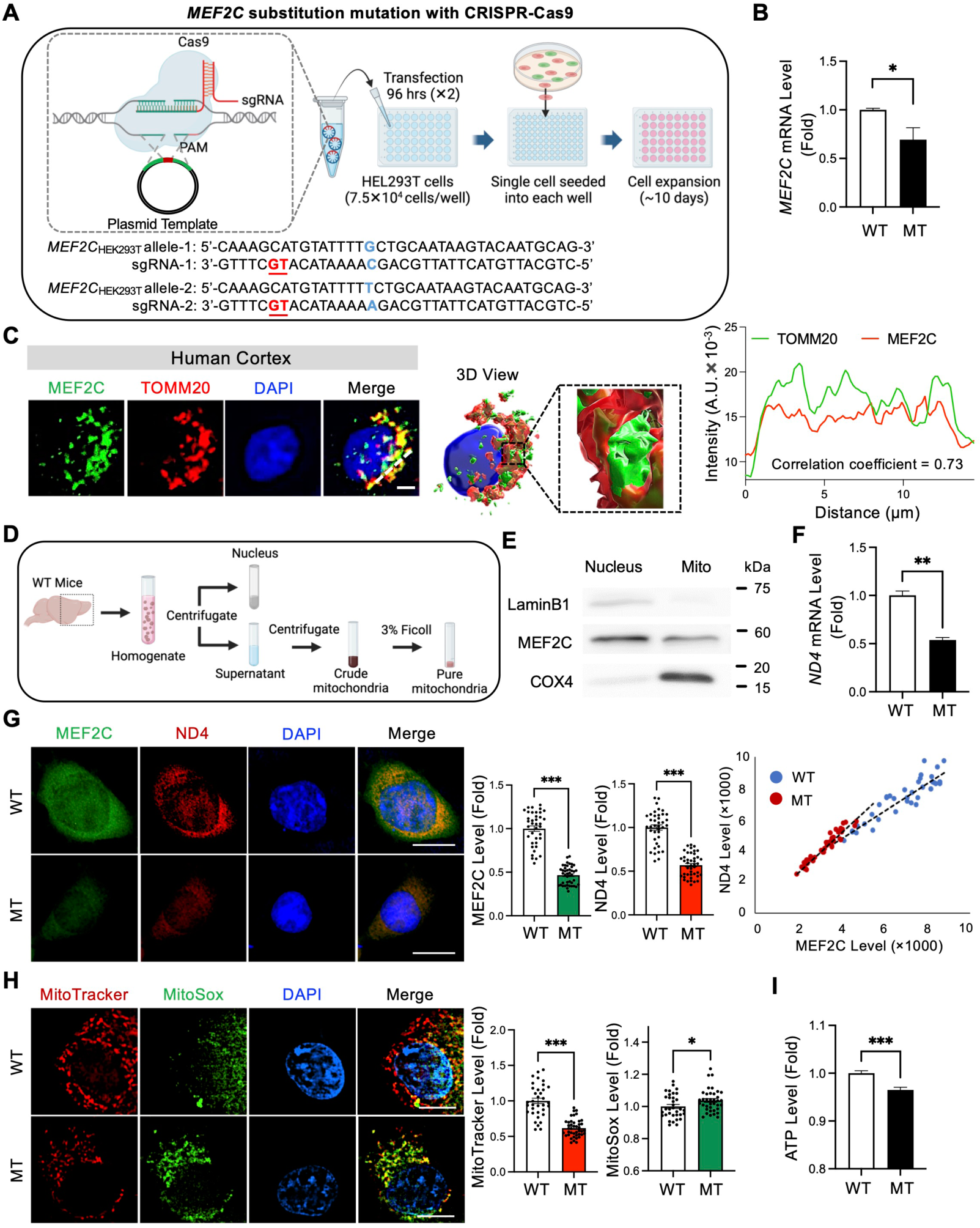
*MEF2C* enhancer mutation impairs MEF2C transcription and leads to mitochondrial dysfunction. **(A)** Demonstration of CRISPR-editing of rs304152-G mutation in HEK293T cells. Scheme created with BioRender.com. **(B)** MEF2C mRNA level decreased in rs304152-G (MT) cells compared to rs304152-T (WT) cells. *\*P* < 0.05; Student’s t test (*n* = 3/group, *df* = 4). **(C)** Immunofluorescence staining of MEF2C (green) and outer mitochondria membrane marker, TOMM20 (red), in human cortical pyramidal neuron along with deconvolved and 3D reconstruction image made by Imaris 9 (Bitplane). Right panel shows line measurement analysis for MEF2C and TOMM20 colocalization signals. White dotted line indicates the colocalization analysis line. **(D)** Ultrafractionation of cellular compartments of mouse brain tissues. **(E)** Western blots of cell fractionation confirmed the presence of MEF2C in mitochondria in pure mitochondrial fractions. **(F)** qPCR results showed decrease of *ND4* mRNA levels in MT cells. *\*\*P* < 0.01; Student’s t test (*n* = 3/group, *df* = 4). **(G)** Immunofluorescence staining of MEF2C and ND4 in WT and MT cells. Scale bars (white): 5 µm. Right: densitometry analysis showed decrease of MEF2C and ND4 levels in MT cells. Scatter plot represents positive correlation between MEF2C and ND4 levels. A total of 90 cells count, 15 cells/well, *N* = 3 (WT, MT). *\*\*\*P* < 0.001; Student’s t test (*n* = 45/group, *df* = 88). **(H)** Immunostaining of MitoTracker (red) and MitoSox (green) in WT and MT cells. The nuclei were counterstained with DAPI (blue). Right: quantitation of MitoTracker and MitoSox levels. A total of 72 cells count, 12 cells/well, *N* = 3 (WT, MT). *\*P* < 0.05, *\*\*\*P* < 0.001; Student’s t test (*n* = 36/group, *df* = 70). **(I)** Decrease of ATP level in MT cells compared to WT cells. The experiment was repeated three times. *\*\*\*P* < 0.001; Student’s t test (*n* = 40/group, *df* = 78). Error bars indicate means ± s.e.m.

We further examined the effect of MEF2C enhancer mutation on mitochondrial-encoded gene transcription and mitochondrial activity. The qPCR results showed the rs304152-G allele had an impaired effect on *ND4* mRNA level (Fig. 3F). The immunofluorescence staining results confirmed the decrease of MEF2C and ND4 immunoreactivity levels in MT cells, and the positive correlation between MEF2C and ND4 immunoreactivity levels (Fig. 3G). The immunofluorescence staining showed less localization of MEF2C in mitochondria and smaller mitochondria network size in MT cells (Supplementary Fig. 6). Immunofluorescence revealed decreased mitochondrial membrane potential and elevated oxidative stress in MT cells (Fig. 3H). Finally, we measured ATP level in cells using luminescence and verified the ATP level significantly decreased in MT cells (Fig. 3I). Altogether, rs304152-G reduced *MEF2C* expression, resulting mitochondria genes dysregulation and mitochondrial dysfunction.

### MEF2C targets mitochondria genome and regulates mitochondria function

After confirming that the *MEF2C* enhancer mutation impaired *MEF2C* and mitochondrial gene transcription, we examined the effect of reduced MEF2C function *in vitro*. Given that MEF2C is a known transcription factor, we investigated the existence of MEF2C binding elements in mitochondria genes. Using the TRANSFAC 6.0-based algorithm Patch 1.0, we predicted the MEF2C-binding sites in mouse mitochondrial DNA (Fig. 4A). To confirm the MEF2C occupancy level in mitochondrial genomes, we performed MEF2C ChIP-qPCR in NSC-34 cells infected by *Mef2c* knock-down (*Mef2c*-KD) virus, pAAV-hSyn(pro)-shMef2c-GFP (Fig. 4B). The *Mef2c*-deficient cells showed reduced Mef2c occupancy in *Nd2*, *Nd4* and *Nd5* genes (Fig. 4C). DNA sequencing of MEF2C-ChIP product verified *Nd2*, *Nd4* and *Nd5* mitochondrial genes (Supplementary Fig. 7A). Moreover, MEF2C occupancy in mitochondrial genomes was confirmed by previous MEF2C-ChIP sequencing data in human umbilical vein cells (GSM809016) and mouse prefrontal cortex (GSM5244364) as well (Supplementary Fig. 7B). To verify whether MEF2C binds to the DNA inside of intact mitochondria or not, we performed nuclear and mitochondria fractionation from the cortex of WT mice. Then, we ran MEF2C-ChIP and qPCR to quantitate MEF2C-DNA occupancy with primers specific to *Nd4*, a mitochondria-encoded gene, and *Rgs6*, a nuclear-encoded gene. *Rgs6* is targeted by MEF2C in cortical neurons of mice (22). As a result, our ChIP and qPCR analysis confirmed that the MEF2C-DNA occupancy in the *Nd4* gene was only found in the mitochondria fraction while the MEF2C-DNA occupancy in the *Rgs6* gene was only detected in the nuclear fraction (Supplementary Fig. 8). *Mef2c*-KD cells exhibited decreased mRNA levels of *Nd2*, *Nd4*, and *Nd5,* while MEF2C overexpression (*MEF2C*-O/E) cells showed increased expression (Fig. 4D-E, Supplementary Fig. 9A-B). We further examined that *Mef2c*-KD significantly decreases *Nd4* mRNA levels in a shMef2c-dose dependent manner. In this context, we verified that *Mef2c*-KD, as a similar experimental model of reduced *Mef2c* expression due to SNP (rs304152), directly and negatively affects mitochondria gene transcription (Supplementary Fig. 9C). MEF2C not only directly regulates mitochondria genes, but also indirectly by regulating nuclear-encoded genes such as mitochondrial transcription factor A (TFAM), translocase of the outer mitochondrial membrane 20 (TOMM20), mitofusin 1 (MFN1) and dynamin-related protein 1 (DRP1) in mRNA expression level (Supplementary Fig. 9D-F). TFAM binds to mitochondrial DNA (mtDNA) and involves in initiating and regulating transcription of mtDNA. TOMM20 is a receptor on the outer mitochondrial membrane involves in the import of precursor proteins into mitochondria, and MFN1 and DRP1 are involved in the balance of mitochondrial fusion and fission. In order to examine whether MEF2C functions as a monomer or dimer in the mitochondria, we crosslinked the mitochondria fraction with 1% glutaraldehyde and ran Western blot analysis. As a result, we found that both monomer and dimer of MEF2C signals were detected in the mitochondria fraction as well as in the nucleus fraction (Supplementary Fig. 5E).

**Figure 4.**
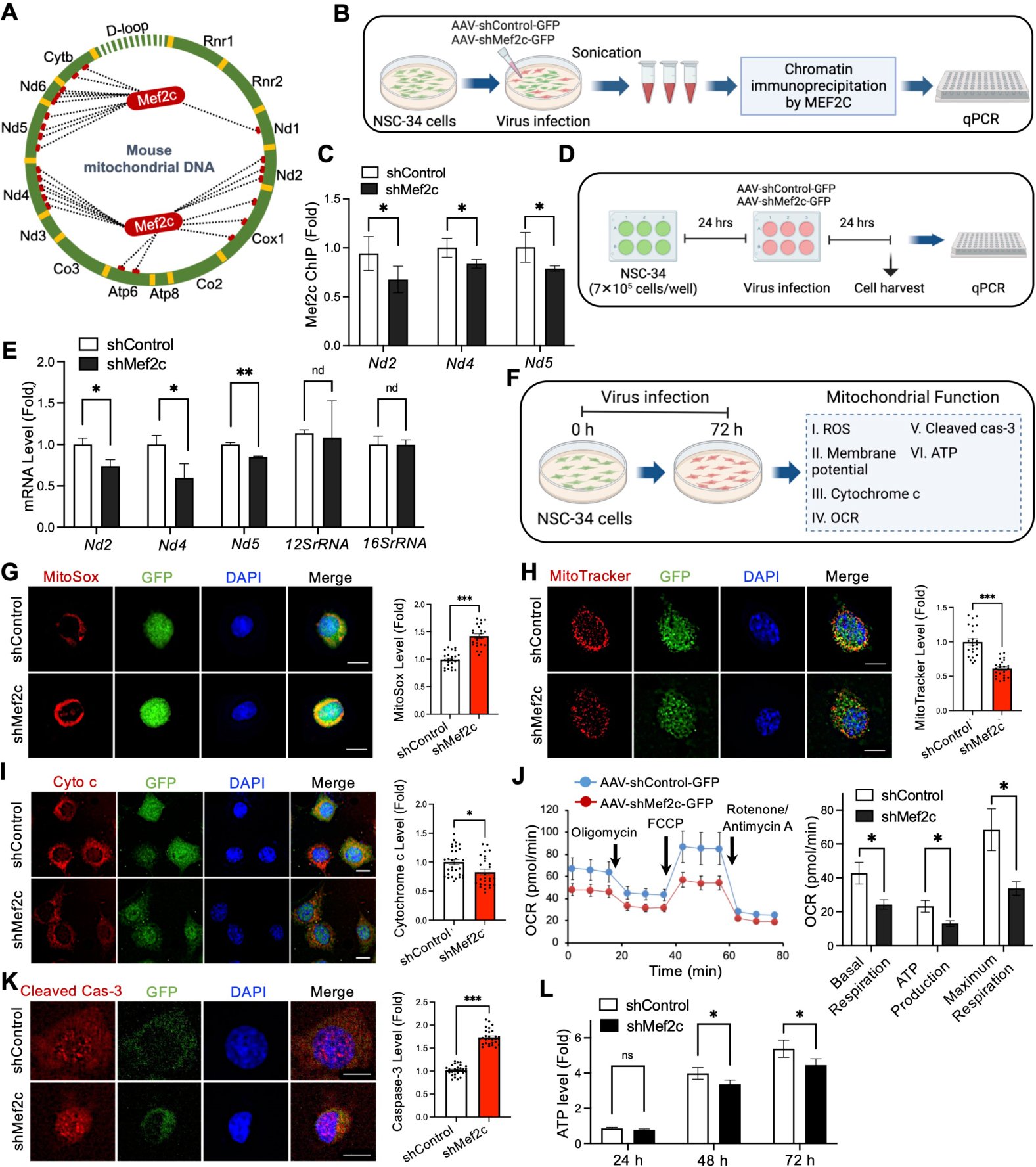
MEF2C regulates mitochondria gene expression and mitochondrial function in motor neuron cell line (NSC-34). **(A)** MEF2C transcription factor binding sites at mouse mitochondrial DNA detected by TRANSFAC 6.0-based algorithm, Patch 1.0. **(B)** A scheme illustrating performing MEF2C ChIP assay on NSC-34 cells infected by *Mef2c*-KD (shMef2c) and control (shControl) viruses, and measuring level of specific DNA in ChIP samples in mitochondria genes by qPCR. **(C)** MEF2C ChIP-qPCR showed Mef2c occupancy level in *Nd2*, *Nd4* and *Nd5* mitochondrial genes in shControl and shMef2c cells. Data generated from 3 samples which are duplicated and normalized to input DNA. *\*P* < 0.05; Student’s t test (*n* = 3/group, *df* = 4). **(D)** A scheme illustrating performing qPCR on NSC-34 cells infected by shControl and shMef2c viruses and harvested after 24hrs. **(E)** qPCR results showed mRNA level of mitochondria genes, *Nd2*, *Nd4*, *Nd5*, decreased by *Mef2c*-KD. *\*P* < 0.05; Student’s t test (*n* = 5/group, *df* = 8). **(F)** A scheme illustrating a series of experiments to evaluate mitochondrial function in in NSC-34 cells infected by *Mef2c*-KD virus. Immunostaining of **(G)** MitoSox (red), **(H)** MitoTracker (red) and **(I)** cytochrome c (red) in GFP^+^ NSC-34 cells infected by *Mef2c*-KD virus. The nuclei were counterstained with DAPI (blue). Scale bars (white): 5 µm. Right: quantitation of MitoSox, MitoTracker and cytochrome c levels in GFP^+^ cells. A total of 60 cells count, 10 cells/well, *N* = 3 (shControl, shMef2c). *\*P* < 0.05, *\*\*\*P* < 0.001; Student’s t test (*n* = 30/group, *df* = 58). **(J)** Cellular respiration rate of differentiated NSC-34 cells infected by *Mef2c*-KD virus measured by a Seahorse XFe24 analyzer. Right: quantitative analysis of the basal and maximum respiratory rate, and ATP production amount of GFP^+^ cells. *\*P* < 0.05; Student’s t test (*n* = 4/group, *df* = 6). **(K)** Immunofluorescence staining of cleaved caspase-3 in NSC-34 cells infected by *Mef2c*-KD virus. Scale bars (white): 5 µm. Right: densitometry analysis for GFP^+^ cells. A total of 60 cells count, 10 cells/well, *N* = 3 (shControl, shMef2c). *\*\*\*P* < 0.001; Student’s t test (*n* = 30/group, *df* = 58). **(L)** Decrease of ATP level by *Mef2c*-KD after 24, 48 and 72hrs virus infection in NSC-34 cells. *\*P* < 0.05; Student’s t test (*n* = 4/group, *df* = 6). Error bars indicate means ± s.e.m.

To investigate the consequences of MEF2C deficiency in mitochondrial function, we performed a series of immunocytochemistry to evaluate mitochondrial ROS, membrane potential, cleaved caspase-3 and cytochrome c levels, and oxygen consumption rate seahorse measurement (Fig. 4F). The mitochondrial superoxide (MitoSox) and mitochondrial membrane potential (MitoTracker) levels were analyzed by comparing the fluorescence staining in NSC-34 cells infected by *Mef2c*-KD and *MEF2C*-O/E viruses (Fig. 4G-H and Supplementary Fig. 9G-H). *Mef2c*-KD resulted in an increase in mitochondrial ROS generation and a decrease in mitochondrial membrane potential in NSC-34 cells (Fig. 4G-H). Moreover, the immunocytochemistry results showed a decrease in cytochrome c level in *Mef2c*-KD cells (Fig. 4I). Then, considering the mitochondria as a main subcellular organelle for cellular respiration and energy production, we verified the *Mef2c* deficiency effects on cellular respiration rate by a Seahorse analyzer (23). The results confirmed the significant decrease of basal and maximum respiration and mitochondrial ATP production rates in *Mef2c*-KD cells (Fig. 4J). In addition, by immunofluorescence labeling, we confirmed an increase in active caspase-3 immunoreactivity in *Mef2c*-KD cells, which indicates induce of apoptotic cell death by *Mef2c* deficiency (Fig. 4K). Finally, the cell viability assay verified a reduction in ATP level by 15% and 17% at 48 and 72hrs after *Mef2c*-KD virus infection in NSC-34 cells, respectively, suggesting elevation of cells energy demand (Fig. 4L). Altogether, MEF2C loss of function increases the mitochondrial oxidative stress, and decrease ATP production, leading to the accumulation of cell damage and eventual cell apoptosis.

### *In vivo* knockdown of *Mef2c* induces mitochondria dysfunction, motor neuronal damage, and altered excitability in mice

In order to verify whether *Mef2c* deficiency affects mitochondrial dysfunction in mice, we delivered AAV-sh*Mef2c* virus into the cortical layer V of wild-type mice by bilateral stereotaxic injection (Fig. 5A). Immunofluorescent labeling revealed a decrease in MEF2C immunoreactivity in GFP^+^ cells within cortical layer V pyramidal neurons of shMef2c mice (Supplementary Fig. 10). Immunostaining with anti-ND4 confirmed that *Mef2c*-KD decreased ND4 immunoreactivity level in mice (Fig. 5B). Consistent with the *in vitro* observation of reduced *Mfn1* and increased *Drp1* mRNA levels, suggesting a shift towards excessive mitochondrial fission following *Mef2c*-KD (Supplementary Fig. 9D), *in vivo* immunostaining with anti-DRP1 as the master regulator of mitochondrial fission showed the increase of DRP1 immunoreactivity level in mitochondria of cortical pyramidal neurons in shMef2c mice (Fig. 5C). Colocalization of DRP1 and the mitochondria outer membrane marker in cortical pyramidal neurons indicates execution of the mitochondria fission process in shMef2c mice (24) (Fig. 5D). The mitochondria fragmentation was quantified by measuring mitochondria network size with TOMM20 signals using MiNA plugin in Fiji ImageJ software (Fig. 5E). To confirm our confocal microscopic observations, we further investigated the MEF2C localization to mitochondria and mitochondria size using transmission electron microscopy (TEM). The immunogold labeling results showed gold-labeled particles in the mitochondria of cortical motor neurons in control mice which reduced from neuronal mitochondria of *Mef2c*-deficient mice (Fig. 5F). Control staining without the primary antibody lacked gold particles within the mitochondria (data not shown). Moreover, TEM images showed mitochondria fragmentation in cortical pyramidal neurons of shMef2c mice (Fig. 5G). Finally, the immunohistochemistry with anti-CTIP2 antibody as a marker for pyramidal neurons layer V showed the shrinkage of pyramidal neurons in shMef2c mice, indicating upper motor neuron damage (Fig. 5H-I) (25). Next, we performed intrathecal injection of *Mef2c*-KD virus in mice and examined the lower motor neuron pathology (Supplementary Fig. 11A). Immunofluorescent labeling confirmed a decrease in MEF2C immunoreactivity levels in GFP^+^ cells in shMef2c mice (Supplementary Fig. 11B). The lower motor neuron *Mef2c*-deficient mice showed a decrease in ND4 but an increase in DRP1 immunoreactivity levels (Supplementary Fig. 11C-E). The marked reduction of ventral neuronal size can be seen in Nissl-stained tissue sections from the lumbar spinal cord of *Mef2c*-KD mice (Supplementary Fig. 11F). Altogether, *Mef2c* deficiency in both upper and lower motor neurons leads to the downregulation of mitochondria genes, mitochondria fragmentation and motor neuronal damage in mice.

**Figure 5.**
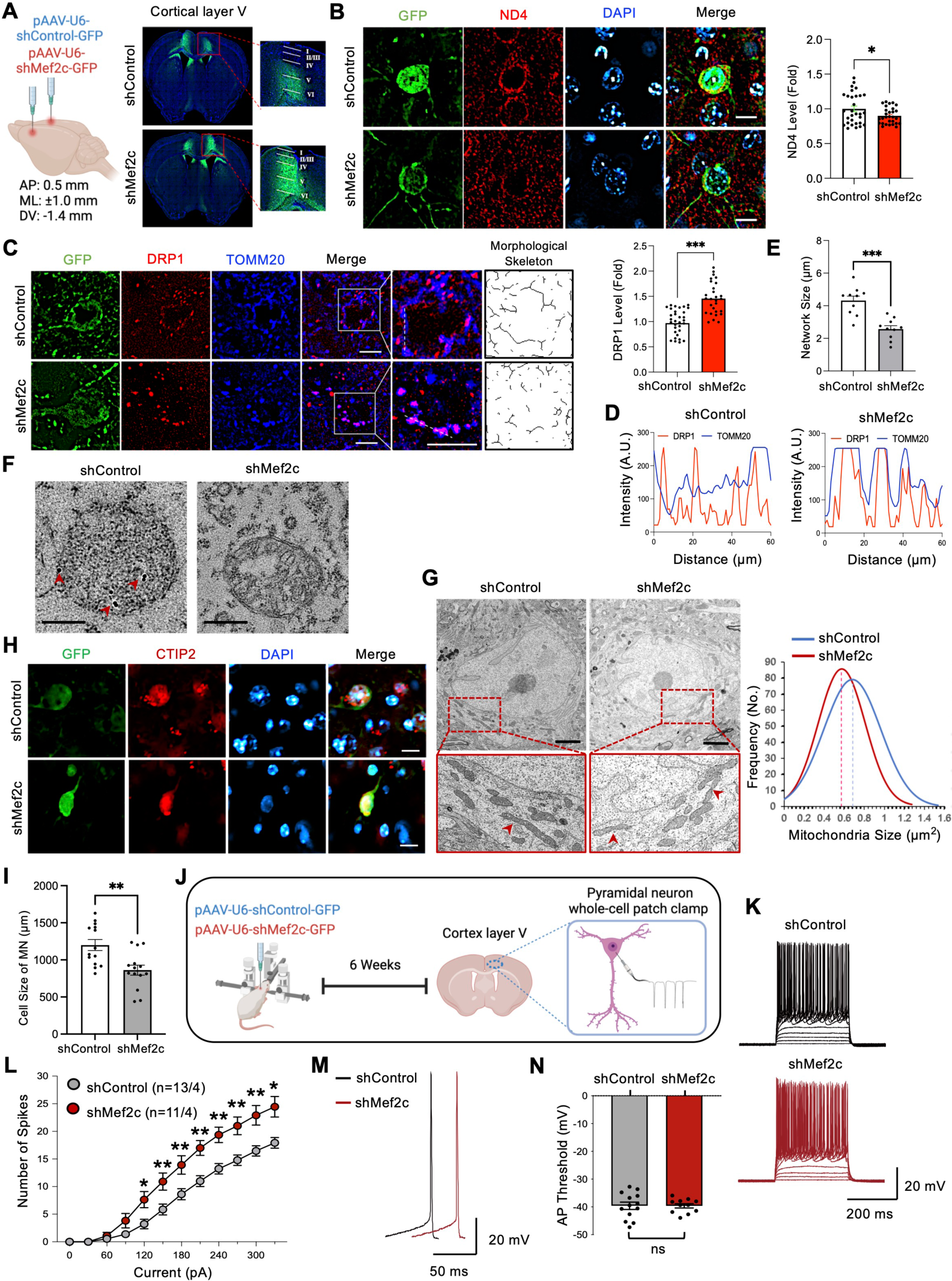
*Mef2c*-KD induces mitochondrial dysfunction and motor neuronal damage, and alters motor neuron excitability in the cortical layer V of mice. **(A)** The pAAV-hSyn(pro)-shControl-GFP (shControl) or pAAV-hSyn(pro)-shMef2c-GFP (shMef2c) viruses was delivered into cortical layer V by bilateral stereotaxic injection method. **(B)** Immunostaining of ND4, GFP, and DAPI in shControl and shMef2c mice. Scale bars (white): 5 µm. Right: densitometry analysis of ND4 immunoreactivity level in GFP^+^ cells. A total of 60 cells count, 6 cells/mouse, *N* = 5 (shControl, shMef2c). *\*P* < 0.05; Student’s t test (*n* = 30/group, *df* = 58). **(C)** Immunostaining of DRP1, TOMM20, and GFP in shControl and shMef2c mice. Scale bars (white): 5 µm. Right: skeletonized images of mitochondria morphological structure using TOMM20 signals, and densitometry analysis of DRP1 immunoreactivity level in mitochondria of GFP^+^ cells. A total of 60 cells count, 6 cells/mouse, *N* = 5 (shControl, shMef2c). *\*\*\*P* < 0.001; Student’s t test (*n* = 30/group, *df* = 58). **(D)** Line measurement analysis for DRP1 and TOMM20 colocalization signals in shControl and shMef2c cortical pyramidal neuron. **(E)** Results of mitochondria network size analysis performed by MiNA plugin in Fiji ImageJ software in 10 cells from each group. *\*\*\*P* < 0.001; Student’s t test. **(F)** Representative TEM images showed the presence of immunogold particles (red head arrows) in mitochondria of pyramidal neurons layer V in shControl mice. Scale bars: 200 nm. **(G)** Representative TEM images showed mitochondria fission in shMef2c group. Scale bars: 2 μm. Right: normal curve for quantitative analysis of mitochondria size in shControl and shMef2c mice cortical pyramidal neurons. *N* = 400 mitochondria/20 neurons (shControl, shMef2c). **(H)** Immunostaining of CTIP2 in shControl and shMef2c mice. Scale bars: 5 µm. **(I)** Quantitation analysis on the cell size of cortical layer V pyramidal neurons in GFP^+^ cells. A total of 30 cells count, 3 cells/mouse, *N* = 5 (shControl, shMef2c). *\*\*P* < 0.01; Student’s t test (*n* = 15/group, *df* = 28). **(J)** A scheme illustrating performing whole-cell patch clamp on cortical layer V pyramidal neurons for shControl and shMef2c mice. **(K)** Representative recordings of action potentials from motor cortex layer V pyramidal neurons induced by step current injection, ranging from 0 to 330 pA (Scale bar: 20 mV, 200 ms). **(L)** shMef2c increased the number of action potentials triggered by a series of current injection (30-pA increment, 12 steps, shControl: n=13 neurons/4 animals, shMef2c: n=11/4). *\*P* < 0.05, *\*\*P* < 0.01; Repeated measures ANOVA test (*df* = 220). **(M)** Representative traces of individual action potential (Scale bar: 20 mV, 50 ms). **(N)** The threshold to initiate action potentials is not altered in shMef2c group (shControl: n=13/4, shMef2c: n=11/4). Student’s t test (*df* = 22). Error bars indicate means ± s.e.m.

Finally, we examined a potential change in intrinsic excitability in upper motor neurons at cortical layer V of *Mef2c*-KD mice (Fig. 5J). Electrophysiological analysis showed that the current step triggered significantly higher number of action potentials in the shMef2c group, indicating that knockdown of *Mef2c* increases the intrinsic neuronal excitability in motor cortex layer V pyramidal neurons (Fig. 5K-L). However, the threshold for firing action potentials was not altered by *Mef2c*-KD (Fig. 5M-N).

### *Mef2c*-KD exhibits motor neuron disease-like behaviors in mice

Finally, we examined whether knockdown of *Mef2c* in upper motor neurons and lower motor neurons affects the motor behavior in mice. The longitudinal behavioral study including open field, tail suspension, accelerated wheel, and inverted grid tests, was performed 3, 6 and 9 weeks after delivering *Mef2c*-KD virus into the cortical layer V of mice by bilateral stereotaxic injection method (Fig. 6A). *Mef2c*-KD mice showed abnormal hindlimb extension reflex compared to control mice (Fig. 6B and Supplementary Fig. 12A). The frequency of forelimb movements was higher in shMef2c mice at 6 and 9 weeks after injection (Fig. 6C). Hindlimb posture is visually reflected by temporally aggregated coordinate plots for the first 10 seconds of tail suspension (Fig. 6D). The inverted grid test was performed as a method to assess muscle strength using all four limbs. The results showed less minimal holding impulse time (body mass × cling time) for *Mef2c*-KD mice, though not statistically significant, a trend towards reduced holding time was observed (Fig. 6E). The computer-assisted footprint analysis allows us to calculate gait characteristics while the mice walk inside of an accelerating wheel (Fig. 6F and Supplementary Fig. 12B). Quantitative analysis of footprint patterns revealed motor coordination impairment in shMef2c mice from 6 weeks post-injection, by wider stride width and shorter stride length (Fig. 6G). Open field test results showed normal behavior in locomotion and anxiety levels in shMef2c mice (Supplementary Fig. 13).

**Figure 6.**
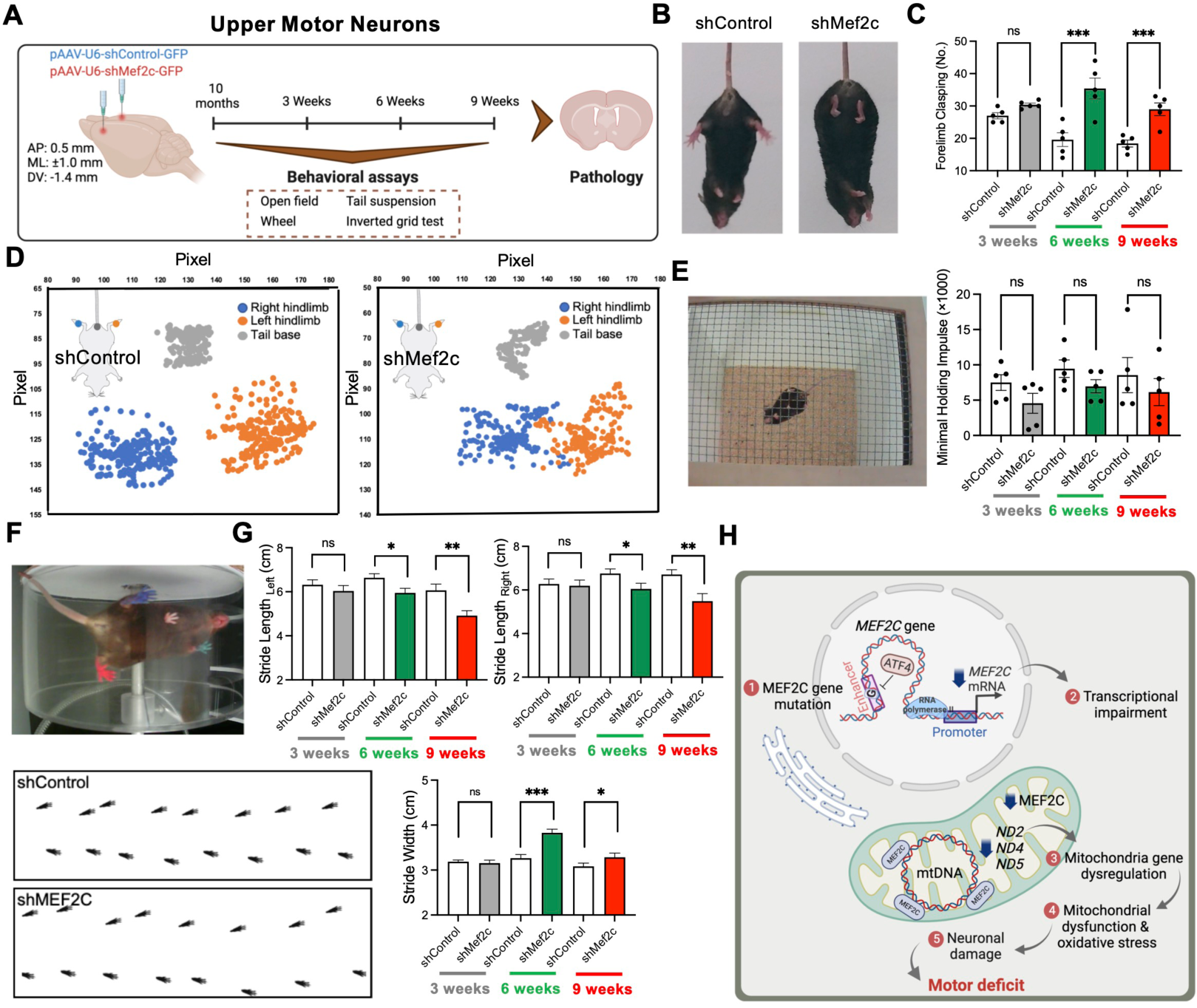
*Mef2c*-KD shows motor neuron disease-like behaviors in mice. **(A)** A scheme illustrating delivery of AAV-shRNA-Control-GFP (shControl) or AAV-shRNA-Mef2c-GFP (shMef2c) viruses into the cortical layer V of WT mice and performing longitudinal behavioral study 3, 6 and 9 weeks after injection. **(B)** Still images of tail suspension of shControl and shMef2c representative mice. **(C)** The number of forelimbs clasping in tail suspension test in 3-, 6- and 9-weeks post-injection. *\*\*\*P* < 0.001; Repeated measures ANOVA test (*n* = 5/group, *df* = 24). **(D)** Aggregated coordinate plots in the first 10 seconds of tail suspension test for shControl and shMef2c representative mice. **(E)** Still image of inverted grid test. Right: the minimal holding impulse. Repeated measures ANOVA test (*n* = 5/group, *df* = 24). **(F)** Still image of accelerated wheel test. Bottom: the computer-assisted footprint in wheel running test for a representative mouse in each group. **(G)** Gait analysis showed shorter stride length and wider stride width in shMef2c mice. *N* = 5 (shControl, shMef2c). *\*P* < 0.05, *\*\*P* < 0.01, *\*\*\*P* < 0.001; Repeated measures ANOVA test (*n* = 30/group, *df* = 116). Error bars indicate means ± s.e.m. **(H)** A scheme proposing how *MEF2C* enhancer SNP impairs *MEF2C* transcription, resulting mitochondrial dysfunction and inducing oxidative stress which makes motor neuronal damage and motor deficit. Scheme created with BioRender.com.

We also examined the motor behavioral changes in lower motor neurons *Mef2c*-KD mice by performing behavioral tests 3, 6 and 9 weeks after intrathecal virus injection (Supplementary Fig. 14A). The *Mef2c*-KD mice showed abnormal hindlimb clasping and increase of number of forelimb clasping (Supplementary Fig. 14B-D). The inverted grid test showed less minimal holding impulse time in *Mef2c*-KD mice (Supplementary Fig. 14E). The computer-assisted gait analysis revealed significantly decrease of stride length and an increase in stride width in *Mef2c*-KD mice hindlimbs (Supplementary Fig. 14F). Furthermore, *Mef2c*-KD mice showed fewer spontaneous movements, specifically decrease of the number of rearing, in the cylinder test at 3- and 6-weeks post-injection (Supplementary Fig. 14G). The behavior tests result at 9-weeks post-injection showed no significant differences between shControl and shMef2c mice because of render inoperative of intrathecal AAV-shRNA virus injection (26) (Supplementary Fig. 15). Open field test results showed normal behavior in locomotion and anxiety levels in lower motor neuron *Mef2c*-KD mice (Supplementary Fig. 16). Altogether, the motor behavior tests demonstrated motor neuron disease-like behaviors in both upper and lower motor neurons *Mef2c*-deficient mice.

We summarize our findings that *MEF2C* enhancer ALS-associated SNP (rs304152) impairs *MEF2C* transcription by inhibiting ATF4 transcription factor binding. Downregulation of *MEF2C* directly dysregulates mitochondria-encoded genes transcription. Finally, MEF2C deficiency leads to mitochondrial dysfunction, oxidative stress and consequently motor neuronal damage, which is the main reason for ALS pathogenesis (Fig. 6H).

## Discussion

This study identifies a novel ALS-associated intronic SNP (rs304152) located in active regulatory regions marked by H3K27ac in humans using a CNN model (27). The rs304152 is located in the enhancer region of the MEF2C gene. The minor allele (T/G) of rs304152 impaired ATF4 binding to *MEF2C* enhancer region, resulting a reduction of the transcriptional activity in its own gene expression (28). Enhancer regions have a profound influence on the regulation of gene expression. The enhancer genetic mutation disrupts the interaction between the enhancer and transcription factor and effects promoter activity (29). Our data indicates, for the first time, that the rs304152 SNP located in the intronic enhancer region of *MEF2C* affects its own gene expression via an epigenetic regulatory pathway of gene transcription.

MEF2C, a neuron-specific transcription factor, is known to be localized to the nucleus and plays a crucial role in the regulation of gene expression (30). Interestingly, in the current study, we discovered mitochondrial localization of MEF2C in cortical pyramidal neurons in human and mouse brains. We verified that MEF2C directly regulates mitochondria genes and mitochondrial function both *in vitro* and *in vivo*. Given that a previous study identified MEF2C-MEF2D heterodimers in HEK293 cells, our results indicate that MEF2C may function as a monomer or dimer in the mitochondria as in the nucleus (31). It is interesting to note that the heterodimer of more than one MEF2 protein does not exhibit better transcriptional activity than the homodimer of a single MEF2 protein (32). In this context, either monomer or dimer of MEF2C is more transcriptionally active in mitochondrial gene regulation, needs to be determined in future studies. Mitochondrial dysfunction has been implicated in various neurodegenerative diseases (33). Previous work has shown that TFAM links the nuclear transcription response to mtDNA content and has a role in supporting mitochondrial respiratory function (34). TOMM20 is a protein that plays a crucial role in mitochondrial respiration, ATP production, and the transport of proteins into mitochondria (35). We determined that the knockdown of *Mef2c* decreases mitochondrial membrane potential and respiration function, leading to a loss of ATP production. In turn, *Mef2c* deficiency increased mitochondrial oxidative stress and induced neuronal apoptosis in NSC-34 motor neurons (36). Furthermore, *Mef2c*-KD mice showed reduction of mitochondria gene expression and excessive mitochondrial fission by the translocation of Drp1 from cytosol to the mitochondrial outer membrane in both upper motor neurons (pyramidal neurons at the cortical layer V) and lower motor neurons (motor neurons at the lumbar spinal cord). Our results suggest that the loss of MEF2C function dysregulates the expression of mitochondria-encoded genes (*ND2*, *ND4* and *ND5*), alters mitochondrial morphology, and induces mitochondrial dysfunction, leading to motor neuronal damage similar to the pathology found in ALS mice (37).

A previous study by Mitchell *et al.* has demonstrated that MEF2C deficiency is a genetic and epigenetic risk factor for SCZ (38). Rescue of Mef2c function improves cognitive performance in working memory and object recognition in mice (38). On the other hand, a recent report shows that the decrease of MEF2C-mediated neuronal transcription by the activation of microglial cyclic GMP–AMP synthase and type I interferon (IFN-I) signaling declines cognitive function in AD (39). In addition, Li et. al. found a de novo autism-associated mutation in the *MEF2C* gene and constructed *Mef2c*-mutant mice showing autistic-like behaviors that can be rescued by CRISPR-Cas9 genome editing (40). However, it is not known whether there is a casual relationship between the enhancer mutation of *MEF2C* gene and the onset of brain disorders. Currently, it is well accepted that disruption of enhancer function through genetic mutation or chromosomal rearrangement is closely linked to disease-driving mechanisms (41). Since we verified that the enhancer mutation of *MEF2C* down-regulates its own gene expression, we utilized *Mef2c*-KD mouse model to investigate the loss of *Mef2c* function *in vivo*. Interestingly, *Mef2c*-KD mice exhibited dystonia-like phenotype and reduction of hindlimb and forelimb muscle strength. Moreover, the longitudinal behavior analyses showed a progressive dysfunction of motor coordination in *Mef2c*-KD mice that is consistent with the behavioral phenotypes of ALS mice (42–45). Building upon previous findings and our own data, we propose that loss of MEF2C function by its enhancer mutation can be a risk factor for motor neuronal dysfunction in mice.

Previous studies have shown that ALS patients and ALS mice (G93A) exhibit hyperexcitability of cortical motor neurons, but the mechanism of hyperexcitability is not known (46, 47). Our electrophysiology data shows a significant increase in intrinsic excitability by increasing the number of action potentials in pyramidal neurons at cortical layer V of *Mef2c*-deficient mice without affecting the threshold for firing action potentials. Consistent with our data, Wainger *et al.* reported hyperexcitability of SOD1 ALS-derived motor neurons and no changes in the threshold of action potentials (46). Once the higher frequency of firing leads to accumulation of Na^+^, Ca^2+^, and Cl^−^ in neuronal cytosol and release of K^+^ to the extracellular space (48), it can cause excitotoxicity, ultimately leading to neuronal damage (49, 50). Additionally, given that mitochondria activity is pivotal for modulating the excitability of motor neuron, MEF2C deficit-associated mitochondrial dysfunction can lead to a pathological hyperexcitability and motor neuronal damage (51).

In conclusion, we identified an ALS-associated SNP, rs304152, located in the distal enhancer of *MEF2C* and verified its mechanistic role as an epigenetic modulator for the transcription of its own gene. The loss of MEF2C function by the rs304152-G allele describes metabolic disruptions in motor neurons preceding motor neuronal degeneration in the pathogenesis of ALS. The longitudinal behavioral study further indicates that *Mef2c* deficiency causes progressive muscle weakness and impairment of motor coordination skills in mice similar to ALS mice phenotypes. Together, this study proves that the noncoding variant of *MEF2C*, rs304152, is a *bona fide* risk factor of motor neuron disease, and provides an insight on how a noncoding genetic mutation drives epigenetic changes in the pathogenesis of motor neuron disease.

## Materials and methods

### Convolutional neural network training

We used genetic associations from large-scale GWAS meta-analyses for ALS and SCZ. After filtering out SNPs with p-value > 5×10^−4^, each association block was identified on the lead-SNP with the lowest p-values and located 1 Mb apart from each other, including the 30 most significant neighboring SNPs (14). Subsequently, epigenetic features including DHS mapping, histone modifications, target gene functions, and transcription factor binding sites (TFBS) were annotated to each SNP in 450 and 1044 association blocks constructed for ALS and SCZ, respectively (14). Finally, we trained a CNN model with uncertain labeling on the epigenetic features map of the association blocks (15). The input data, comprising all association blocks (labeled 1), and control blocks constructed by shuffling the regulatory features of SNPs in the association blocks (labeled 0), were divided into training, validation, and testing sets by chromosome (15, 52). Subsequently, the CNN model was trained with uncertain labels on the extracted epigenetic feature map using a large number of hyperparameters and an autoencoder for pre-training, as explained in the reference (15).

### Human post-mortem brain samples

Neuropathological processing of normal and ALS postmortem brain samples was performed using procedures previously established by the Boston University Alzheimer’s Disease Center (BUADC, USA). Cortical and spinal cord histopathology was verified and gra ded according to the criteria as described previously (Myers et al., 1988; Vonsattel et al., 1985). This study was approved for exemption by the Institutional Review Board of the BU School of Medicine (Protocol H-28974) as it included post-mortem tissues that were not classified as human subjects. The study was undergone in accordance with institutional regulatory guidelines and the principles of human subject protection outlined in the Declaration of Helsinki. Information about the brain tissues is provided in Supplementary Table 2.

### Luciferase reporter assay

Duplex of the *MEF2C* enhancer sequence containing a mutation site [5′-ATGTATTTTTCTGCAATAAGT-3′ (×2)] in human genomic DNA were synthesized (Invitrogen). Additionally, 300-bp and 500-bp fragments of human *MEF2C* promoter (from -300 and -500 to −1-bp relative to the translation start site) were generated by PCR from human genomic DNA with primers shown in (Supplementary Table 3). The amplified PCR products were subcloned into the TA vector using TOPclonerTM TA core kit (Emzynomics, Korea) and confirmed by sequencing (Macrogen, Korea). The cloned sequences were released by restriction digestion with BglIII and HindIII and subcloned into the pGL4.14-Enhancer-Luc vector (Promega, USA) at the identical sites.

### Enhancer mutation by CRISPR-Cas9

HEK293T cells (ATCC, CRL-11268) were cultured in Dulbecco’s Modified Eagle Medium (Welgene, USA), supplemented with 10% fetal bovine serum (Welgene, USA) and 1% antibiotic-antimycotic solution (Welgene, USA). For transfection, HEK293T cells (Invitrogen, USA) were seeded at 7.5 × 10^4^ cells per well in 48-well plates. After ∼24 hours, cells were transfected at 70–80% confluency with plasmids encoding Cas9-NG (750 ng) and sgRNA (250 ng) using Lipofectamine 2000 (1.5 µL, Invitrogen, USA). After 96 hours of treatment, cells were subjected to a second treatment to increase the editing efficiencies, following the above process. At 96 hours post-double treatment, we isolated single-cell-derived clonal populations using limiting dilutions. CRISPR-treated cells were seeded at 0.5 cells per well of a 96-well plate. Each cell was grown for ∼10 days and maintained as a clonal cell line. Some cells from clonal populations were harvested using 50 µL of cell lysis buffer (50 mM Tris-HCl, pH 8.0 (Sigma-Aldrich), 1 mM EDTA (Sigma-Aldrich, USA), 0.005% sodium dodecyl sulfate (Sigma-Aldrich, USA), supplemented with 2.5 µL of Proteinase K (Qiagen, USA). Cells were lysed by incubation at 55 °C for 1 hour, followed by incubation at 95 °C for 10 minutes. Allele frequencies of each clonal cell were measured using targeted deep sequencing. To generate NGS libraries, we performed nested PCR using KAPA HiFi HotStart DNA polymerase (Roche, USA) to label each fragment with Illumina adapters and index sequences. Final products were purified using a PCR purification kit (Enzynomics, Korea) and sequenced using a MiniSeq sequencer (Illumina, USA). NGS data were analyzed using the program as previously described (53). The PCR primer sequences are shown in Supplementary Table 4.

### Measurement of oxygen consumption rate

The oxygen consumption rate was measured by a Seahorse XFe24 analyzer (Seahorse Bioscience, USA). A total of 4 × 10^4^ differentiated NSC-34 cells were plated in XFe24 cell culture microplates and cultured. The cartridge plate was incubated with XF Calibrant buffer for one day (37 °C, CO^2^-free); analytical medium (XF basal medium supplemented with 1mM pyruvate, 4mM glutamine, and 25mM glucose) was prepared right before analysis.

## Acknowledgements

We thank Prof. Eun Kyung Lee (Catholic University of Korea) for providing MEF2C construct. We also thank Dr. Chaebin Kim for his assistance in establishing of computer-assisted footprint in accelerated wheel test.

## Authors contributions

A.Y-J., J.L., and H.R. conceived and designed the study and wrote the manuscript. S-H.C. and J.C. performed cell experiments and quantitative RT–PCR analysis. K.L. and J.C. prepared CRISPR-Cas9 cells. P.T-T.N., S.K., and U.P. performed mice injection. A.Y-J. P.T-T.N. and U.P. performed the behavioral tests. P.T-T.N., Y.K., U.P. and S.J.H. performed the ICC, IHC. U.P., S.J.H., H.L.R., G.L., T.D.S. and J.L. provided human tissue collections and performed the immunostaining in the human samples. H.R(Rhim)., H.H. performed the electrophysiological experiment. K.L. performed immunogold staining and TEM imaging. S.K., and J.C. generated the AAV viruses. J.L. and H.R. supervised the analysis and manuscript. All authors contributed to the analysis and discussion of the results and reviewed the manuscript.

---

**Supplementary Fig. 1.**
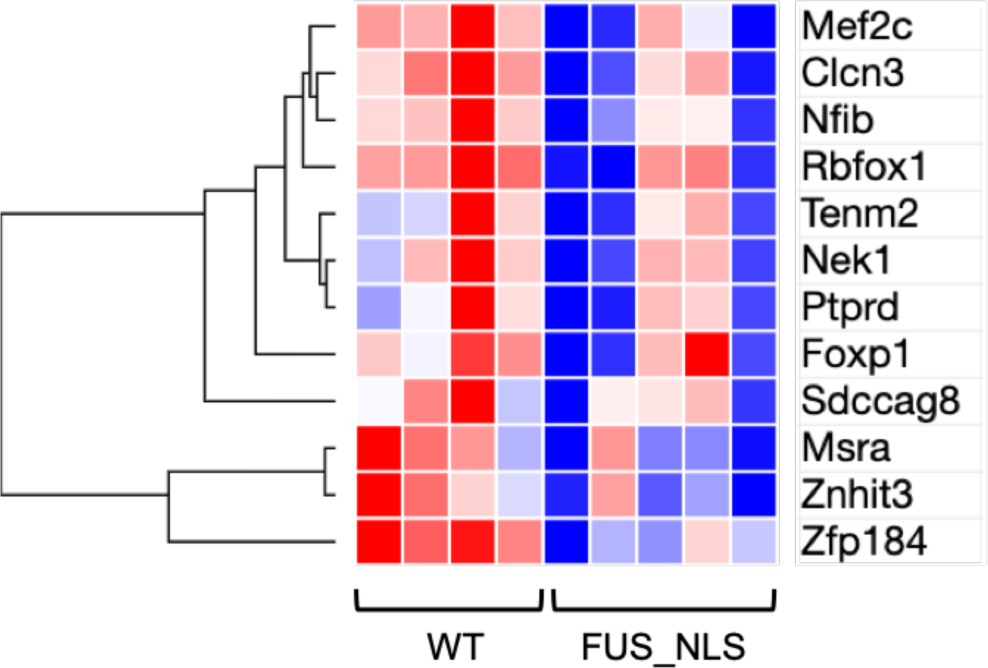
Heatmap analysis for selected 12 genes by CNN algorithm in frontal cortex mRNA transcriptome data from 22-month-old FUS-NLS male mice (19).

**Supplementary Fig. 2.**
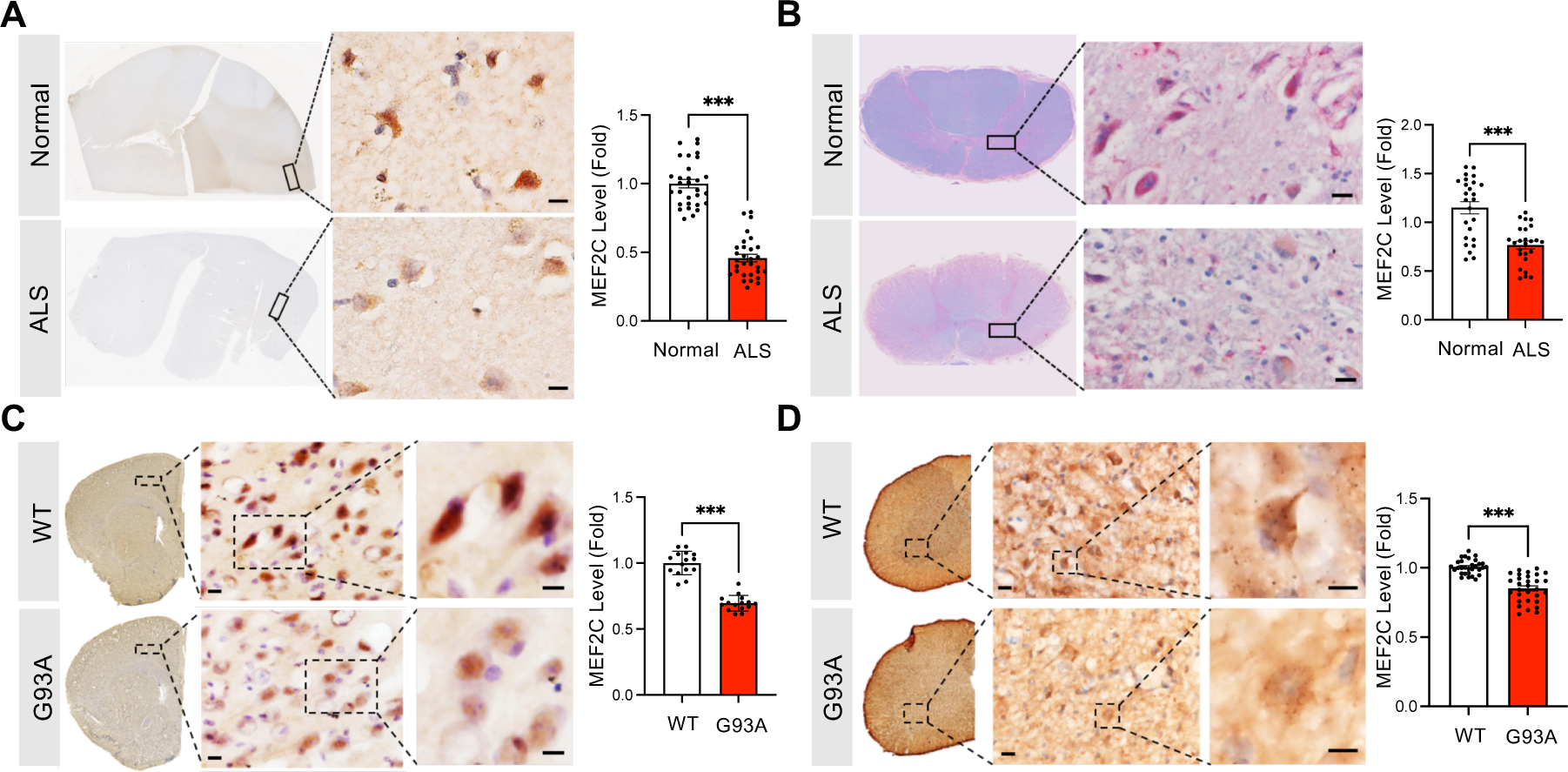
MEF2C immunoreactivity is decreased at cortical layer V pyramidal neurons and spinal cord of ALS patients and ALS mouse model (G93A). Representative DAB staining images show MEF2C immunoreactivity in **(A)** cortical layer V pyramidal neurons. *N* = 5 [Normal, ALS] and **(B)** spinal cord of ALS postmortem and normal subject. Scale bars: 5 µm. *N* = 2 [Normal, ALS]. Right panels show densitometry results of MEF2C immunoreactivity in both cortical layer V pyramidal neurons and spinal cord motor neurons. *\*\*\*P* < 0.001; Student’s t test (*n* = 30, *df* = 58). Representative DAB staining images show MEF2C level in **(C)** cortical layer V pyramidal neurons and **(D)** spinal cord of ALS mouse model (G93A) and control. Right panels show the quantitation of MEF2C immunoreactivity level. Scale bars: 5 µm. *N* = 5 [WT, G93A]. *\*\*\*P* < 0.001; Student’s t test (*n* = 30, *df* = 58). The nuclei were counterstained with hematoxylin. Error bars indicate means ± s.e.m.

**Supplementary Fig. 3.**
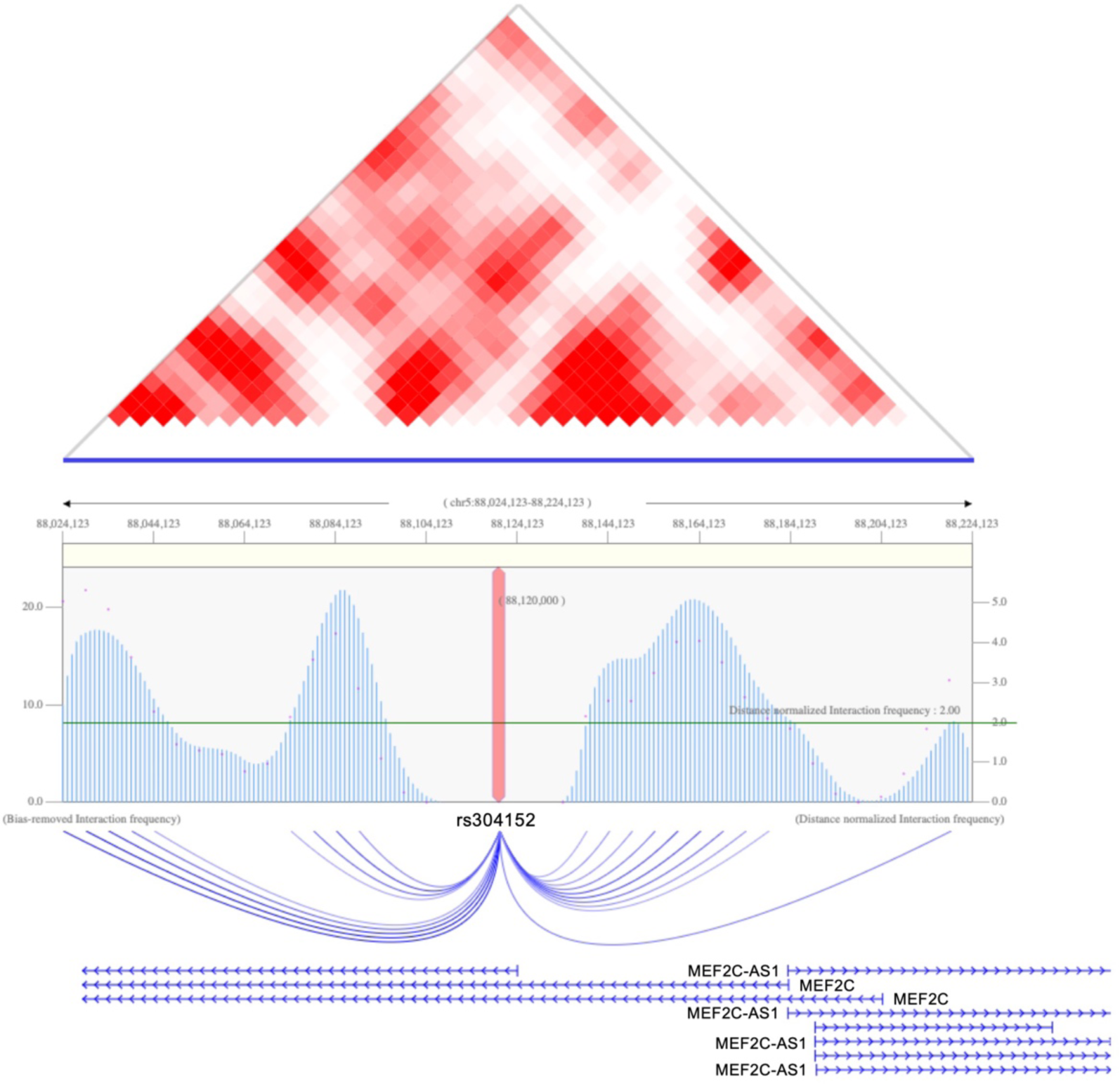
Hi-C data showed the long-range interactions between rs304152 and *MEF2C* promoter in human prefrontal cortex. The TAD result and interaction peaks of rs304152 and nearby genes within a 100 kb region using Hi-C data across human prefrontal cortex. Data from the 3DIV database[14].

**Supplementary Fig. 4.**
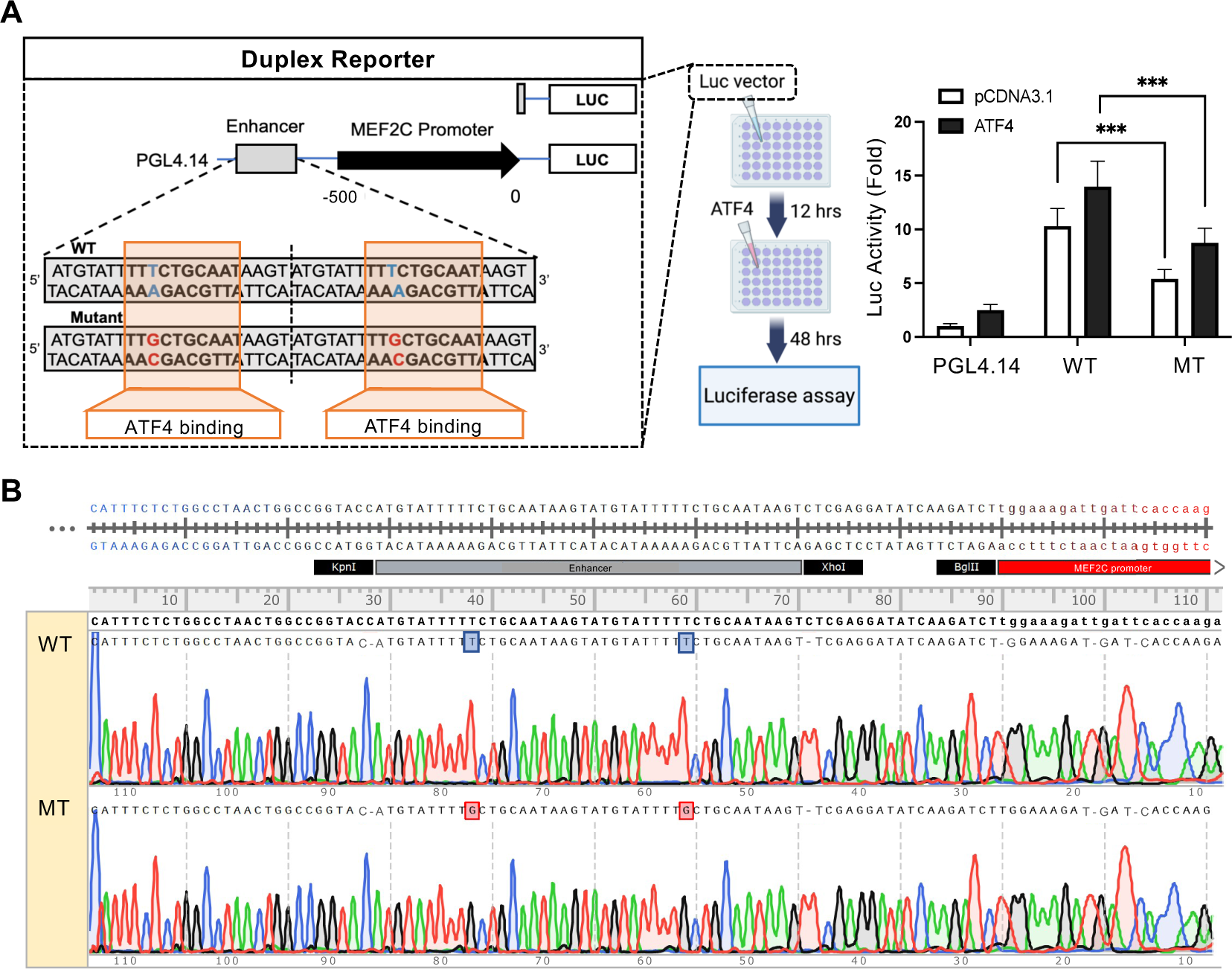
(A) Schematic representation of constructs used in the luciferase reporter assays. Right: The construct containing rs304152-G allele (MT) showed less luciferase activity than the construct containing rs304152-T allele (WT). The experiment was repeated three times (n = 12). *\*\*\*P* < 0.001; Two-way ANOVA test (*n* = 12, *df* = 66). Error bars indicate means ± s.e.m. **(B)** DNA sequencing and chromatograms show that ATF4 ChIP DNA samples from *MEF2C* enhancer region contains duplex of WT or MT form. Alignment was generated by SnapGene software (GSL Biotech; available at snapgene.com).

**Supplementary Fig. 5.**
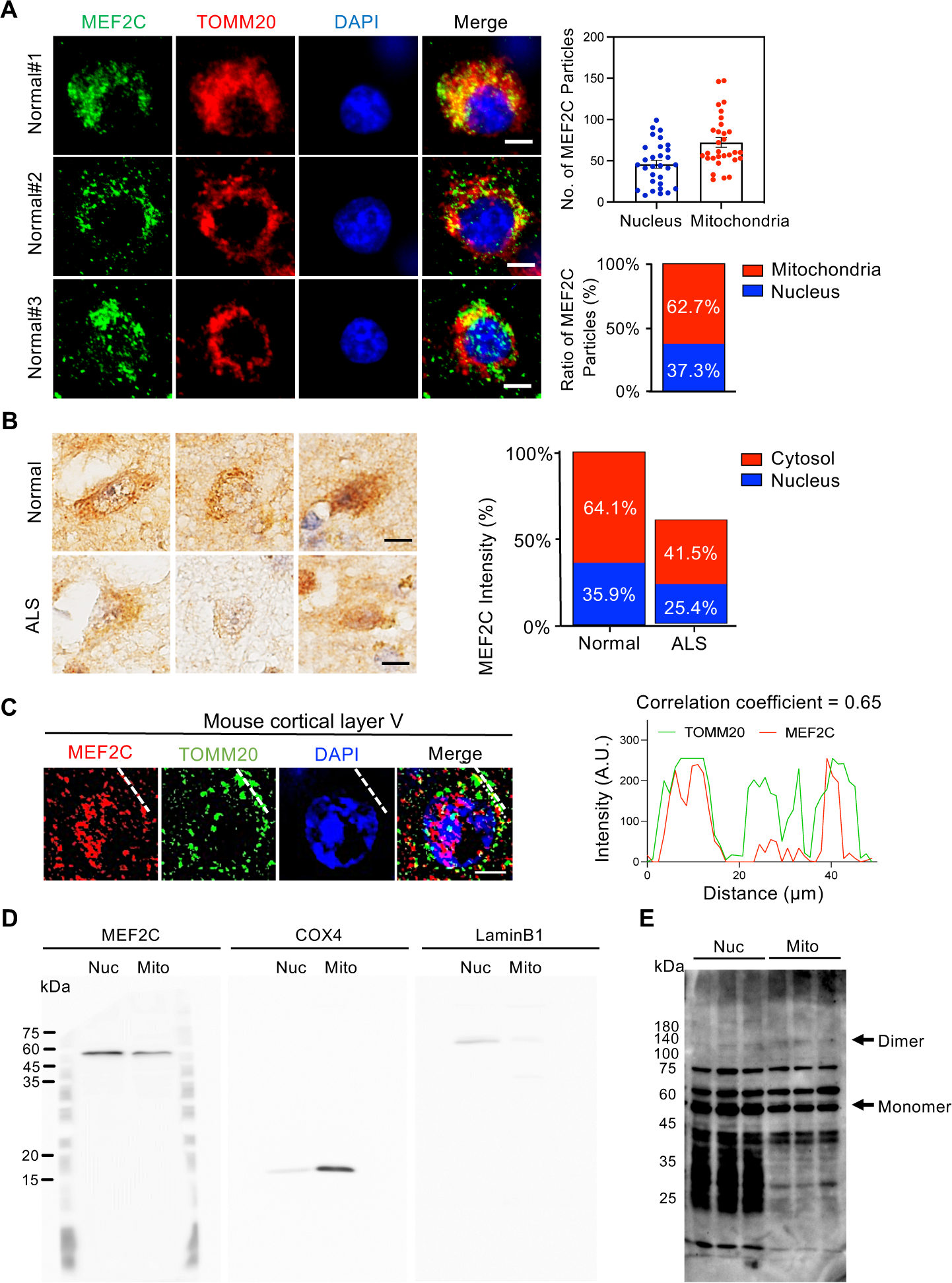
MEF2C localized to the mitochondria of pyramidal neurons in human postmortem brains and mouse motor cortex (layer V). **(A)** Immunofluorescence staining of MEF2C (green) and TOMM20 (red), a mitochondria outer membrane marker, in cortical pyramidal neurons of human postmortem brains. Scale bars (white): 5 µm. Right panels show the number and ratio of MEF2C-positive signals in the nucleus and the mitochondria. A total of 40 cells count, 10 cells/case, *N* = 4 (3 cases are shown above and 1 case is derived from Main Figure 3C). **(B)** DAB staining images present MEF2C immunoreactivity in the cytosol of pyramidal neurons in mouse motor cortex (layer V). The nuclei were counterstained with hematoxylin (blue). Scale bar: 5 µm. Right panel shows the percentile of MEF2C intensity in the nucleus versus the cytosol. A total of 100 cells count, 10 cells/case, *N* = 5 normal control subjects, and *N* = 5 ALS patients. **(C)** Immunofluorescence staining of MEF2C (red) and TOMM20 (green) in the pyramidal neurons of mouse motor cortex (layer V). White dashed line indicates the colocalization analysis foci of MEF2C and TOMM20. Scale bars (white): 5 µm. Right panel exhibits the histogram of MEF2C and TOMM20 colocalization. **(D)** Full blots (developed with MEF2C, COX4 and LaminB1 antibodies) from Western blot analysis for the subcellular fractionation of mouse brain. **(E)** Western blot analysis for detecting MEF2C monomer and dimer from the subcellular fractions crosslinked with 1% glutaraldehyde

**Supplementary Fig. 6.**
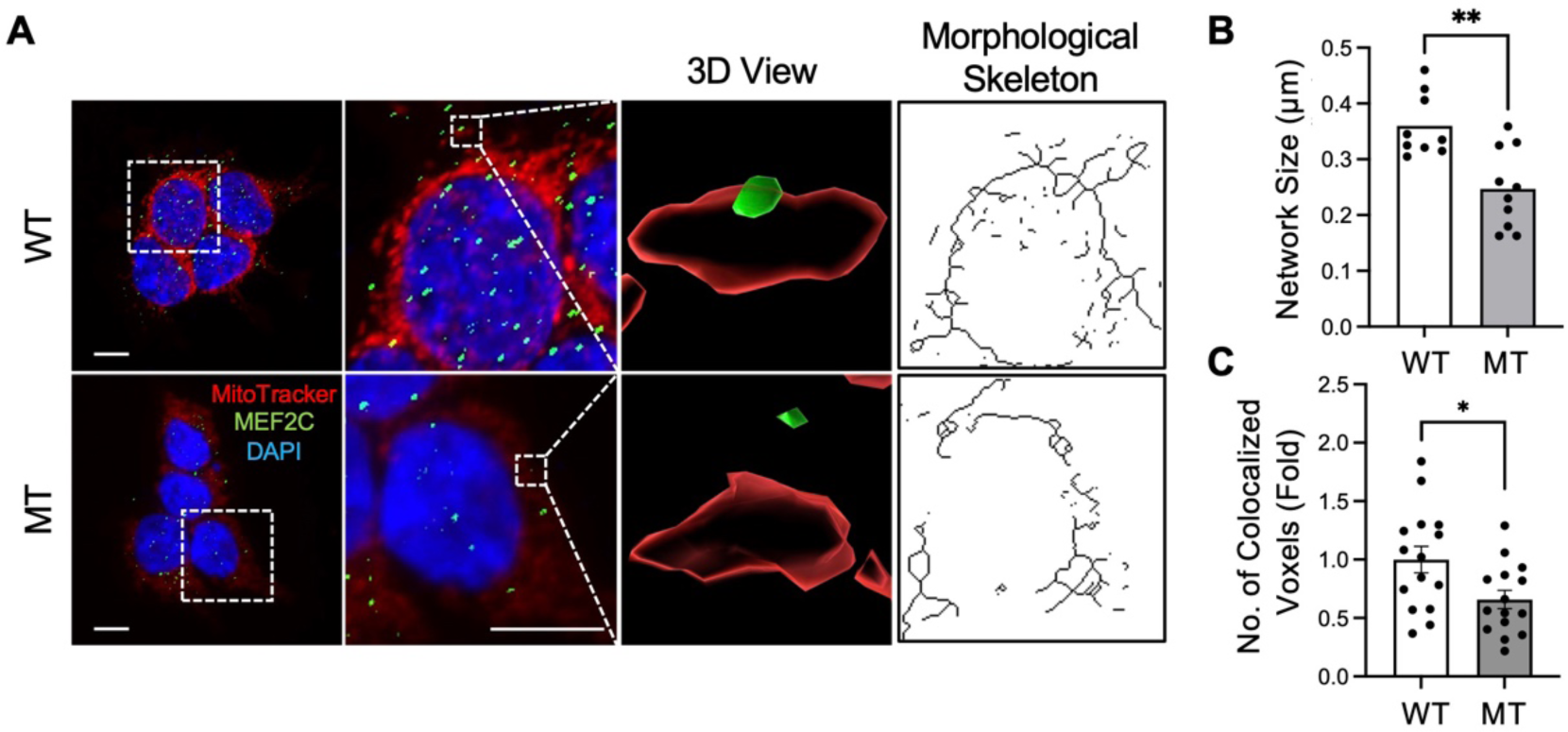
CRISPR-Cas9 cells derived *MEF2C* enhancer mutation exhibits less localization of MEF2C in mitochondria of HEK293T cells. **(A)** Immunofluorescence staining of MEF2C (green) and MitoTracker (red) in CRISPR-Cas9-WT and -MT cells along with 3D reconstruction image made by Imaris 9 (Bitplane). Right: skeletonized images of mitochondria morphological structure by MitoTracker signals. **(B)** Mitochondria network size analysis performed by MiNA plugin in Fiji ImageJ software in 10 cells from each group. *\*\*P* < 0.01; Student’s t test (*n* = 10/group, *df* = 18). **(C)** Quantitation of the number of MEF2C and MitoTracker colocalized voxels made by Imaris 9. *N* = 3 (WT, MT). *\*P* < 0.05; Student’s t test (*n* = 15/group, *df* = 28).

**Supplementary Fig. 7.**
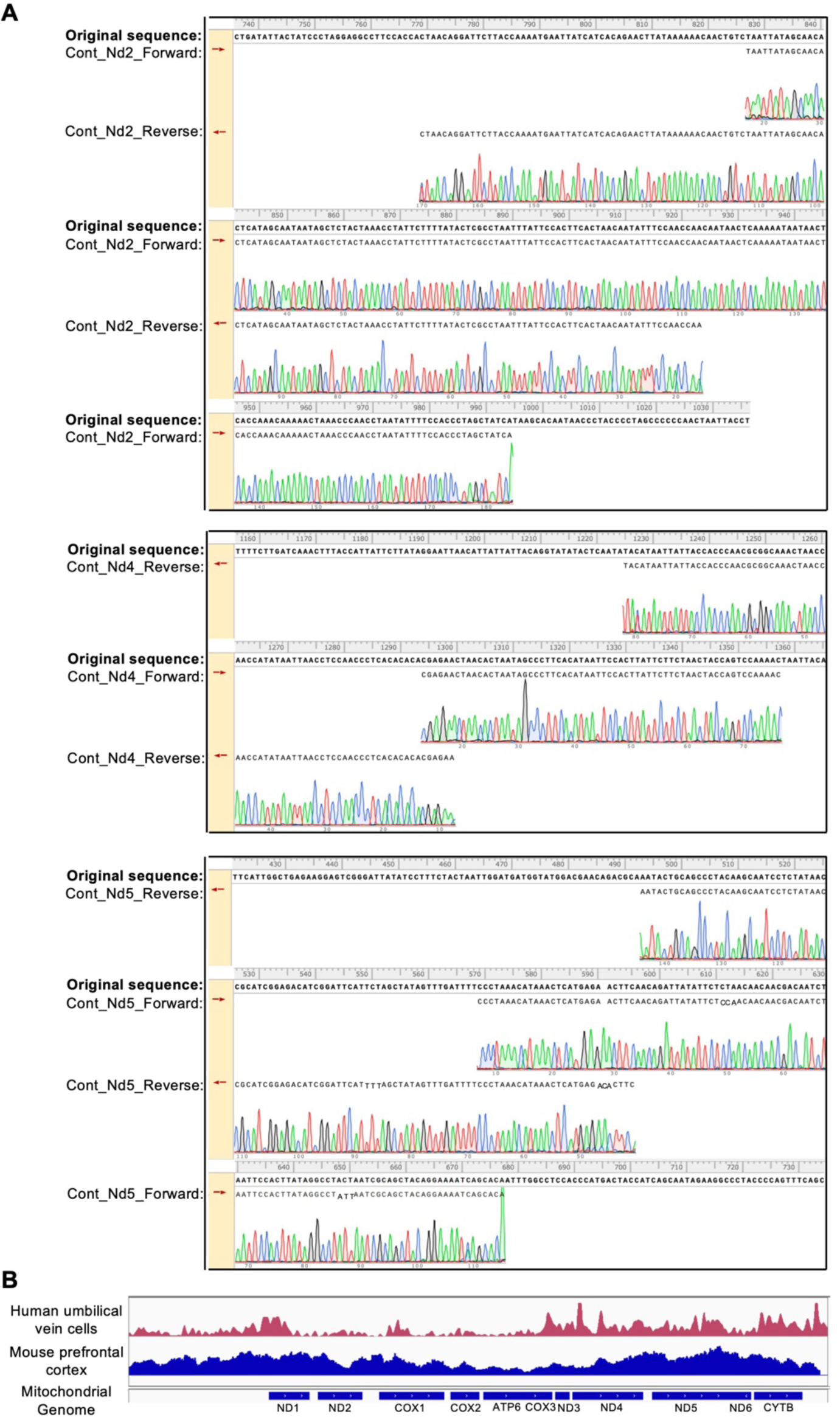
(A) DNA sequencing and chromatograms show that MEF2C-ChIP DNA samples from NSC-34 cell line contain mitochondrial *Nd2*, *ND4* and *ND5*. DNA sequences from MEF2C-ChIP were aligned with mouse mitochondrial genes, and alignment was generated by SnapGene software (GSL Biotech; available at snapgene.com). **(B)** Integrative Genome Viewer (IGV) peak reads of MEF2C in mitochondrial genomes in human umbilical vein cells (GSM809016) and mouse prefrontal cortex (GSM5244364). Alignment was generated by SnapGene software (GSL Biotech; available at snapgene.com).

**Supplementary Fig. 8.**
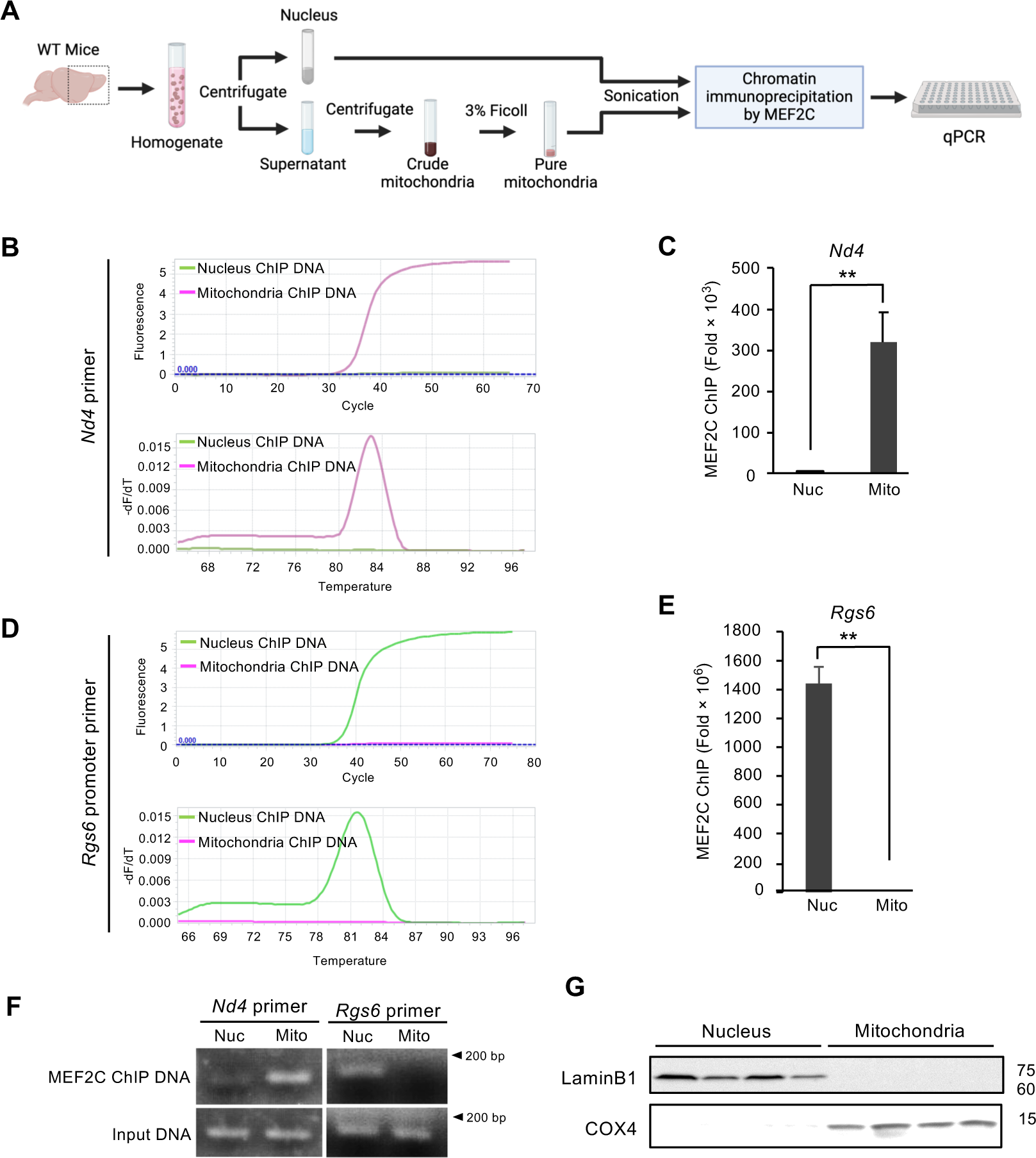
MEF2C binds to mitochondria DNA in the cortex of mouse brains. **(A)** A scheme illustrating the procedure of nuclei and mitochondria fractionation by Ficoll density centrifugation from the cortex of four WT mice. MEF2C ChIP with subcellular fractions and quantification of MEF2C binding to mitochondria-versus nucleus-specific DNA by qPCR were performed sequentially. **(B)** Amplification (ΔCt) (top) and melting (bottom) curves show that DNA eluted from MEF2C ChIP for the mitochondria fraction is amplified with *Nd4*-specific primer but not with *Rgs6*-specific primer. **(C)** MEF2C-DNA occupancy with *Nd4*-specific primer is robustly detected in the mitochondria fraction but not in the nucleus fraction. **(D)** Amplification (ΔCt) (top) and melting (bottom) curves show that DNA eluted from MEF2C ChIP for the nucleus fraction is amplified with *Rgs6*-specific primer but not with *Nd4*-specific primer. **(E)** MEF2C-DNA occupancy with *Rgs6*-specific primer is robustly detected in the nucleus fraction but not in the mitochondria fraction. Data generated from 3 samples that were duplicated. **P < 0.01; Student’s t test (*n* = 3/group, *df* = 4). Error bars indicate mean ± SEM. **(F)** Agarose gel electrophoresis confirmed that MEF2C-DNA occupancy with *Nd4*-specific primer is found in the mitochondria fraction while MEF2C-DNA occupancy with *Rgs6*-specific primer is found in the nucleus fraction. **(G)** Western blot analysis was performed to verify the purity of subcellular fractionations by specific antibodies for COX4, a mitochondria marker, and LaminB1, a nucleus marker (*N* = 4).

**Supplementary Fig. 9.**
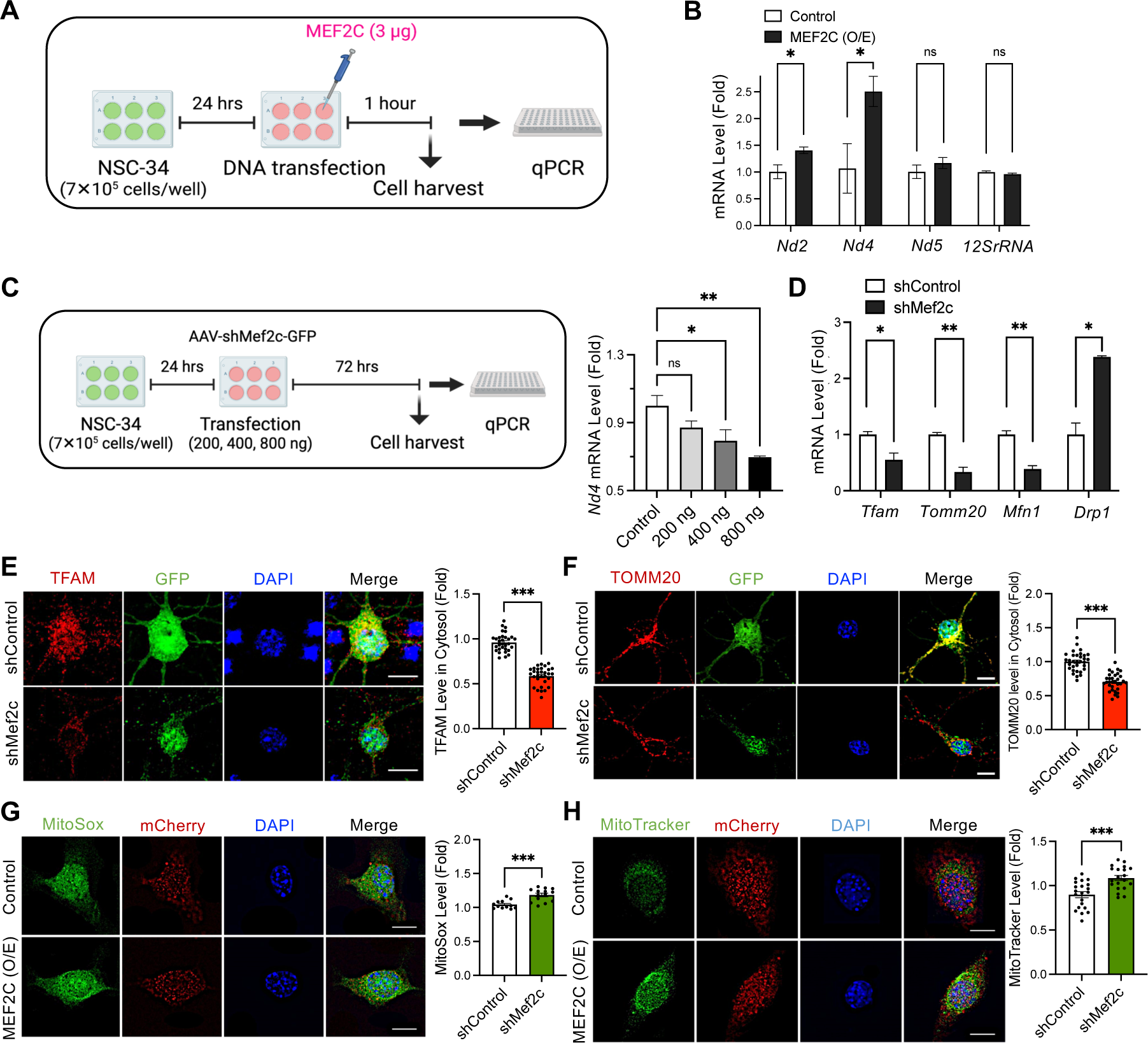
MEF2C regulates nuclear and mitochondria-encoded genes in motor neuron (NSC-34) cells. **(A)** A scheme illustrating MEF2C transfection in NSC-34 cells, RNA isolation and cDNA synthesis from 3 samples. **(B)** qPCR results show mRNA level alteration of mitochondria genes, *Nd2*, *Nd4*, *Nd5* and *12SrRNA* by MEF2C-O/E. *\*P* < 0.05; Student’s t test (*n* = 3/group, *df* = 4). **(C)** A scheme illustrating a dose-dependent shMef2c transfection (200 ng, 400 ng, and 800 ng per well) in NSC-34 cells. The right panel graph shows that *Nd4* mRNA level is reduced by shMef2c in a dose-dependent manner. **(D)** qPCR results show mRNA level of nuclear-encoded genes, *Tfam*, *Tomm20, Mfn1* and *DRP1* in *Mef2c*-KD cells. *\*P* < 0.05, *\*\*P* < 0.01; Student’s t test (*n* = 3/group, *df* = 4). Immunocytochemistry with **(E)** anti-TFAM and **(F)** anti-TOMM20 level in mouse cortical primary neurons infected by *Mef2c*-KD virus for 72 hrs. Right panels are quantitative results of TFAM and TOMM20 levels in the cells cytosol. A total of 60 cells count, 10 cells/well, *N* = 3 (shControl, shMef2c). *\*\*\*P* < 0.001; Student’s t test (*n* = 30/group, *df* = 58). Scale bar (white): 5 μm **(G)** MEF2C-O/E in NSC-34 cells increased both MitoSox (green), and **(H)** MitoTracker signals (green). The nuclei were counterstained with DAPI (blue). Scale bars (white): 5 μm. Right panels are quantitation of MitoSox and MitoTracker level in GFP^+^ cells. *N* = 3 (Control, MEF2C(O/E)). *\*\*\*P* < 0.001; Student’s t test (*n* = 20/group, *df* = 38). Error bars indicate means ± s.e.m.

**Supplementary Fig. 10.**
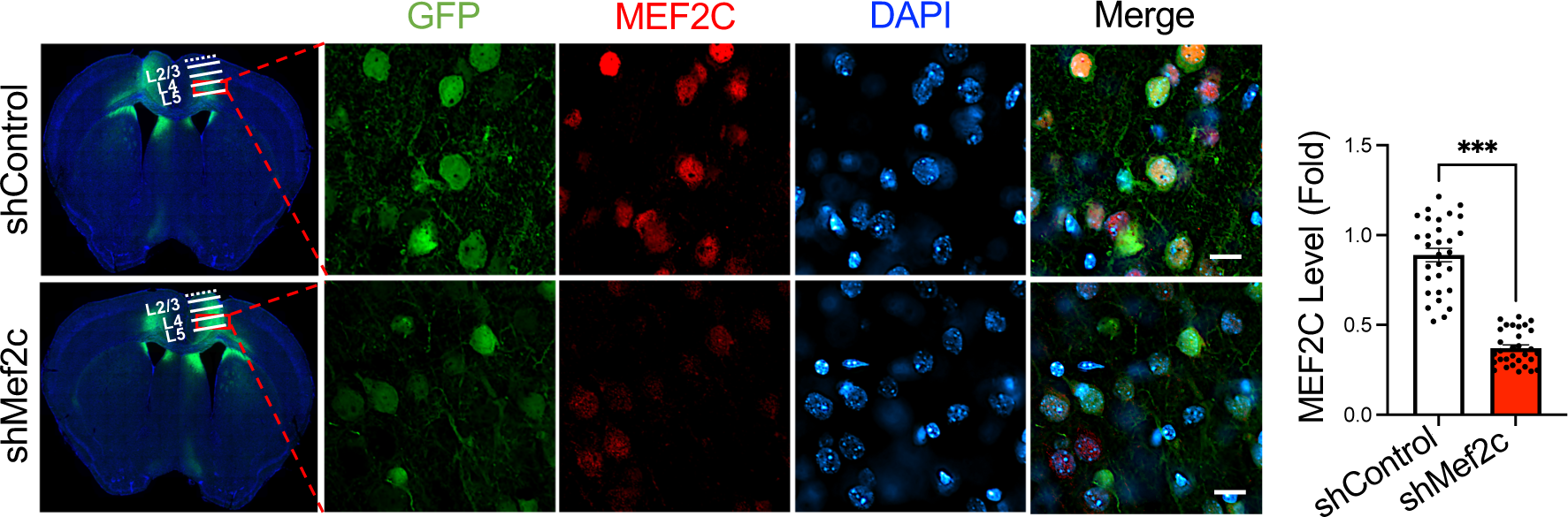
MEF2C level reduced in pyramidal neurons at cortical layer V *Mef2c*-KD mice. The immunostaining illustrated MEF2C level at 9 weeks after delivery of *Mef2c*-KD virus into cortical layer V. Scale bars (white): 5 µm. Right panel shows quantitative results of MEF2C level in GFP^+^ cells. A total of 60 cells count, 6 cells/mouse, *N* = 5 (shControl, shMef2c). *\*\*\*P* < 0.001; Student’s t test (*n* = 30/group, *df* = 58). Error bars indicate means ± s.e.m.

**Supplementary Fig. 11.**
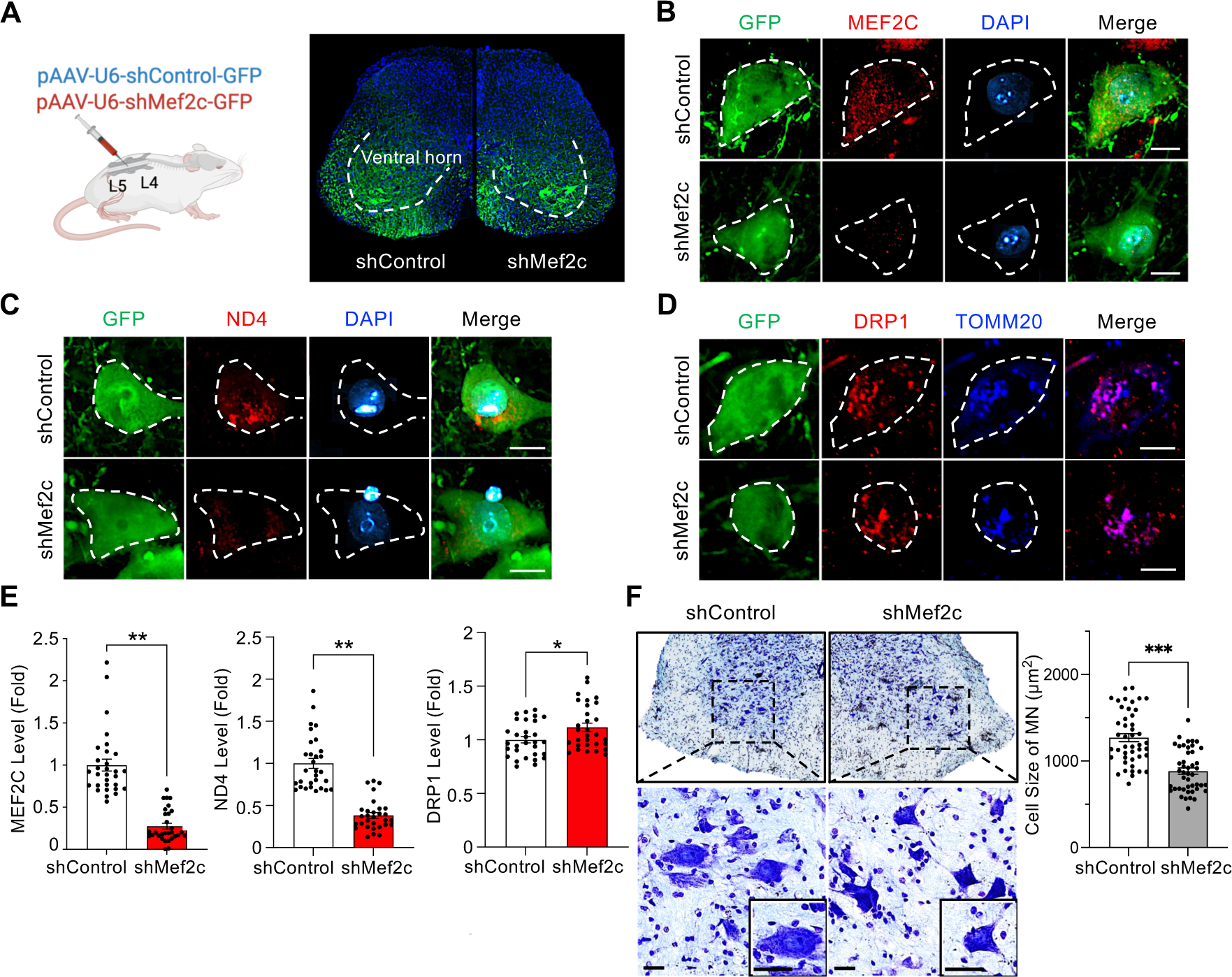
*Mef2c*-KD in the lumbar spinal cord leads to mitochondrial dysfunction and motor neuronal damage in mice. **(A)** A scheme of intrathecal delivery of AAVs in mice (left panel). Detection of GFP and DAPI signals verified that AAV was delivered to the spinal cord ventral horn of mice (right panel). The immunostaining with **(B)** anti-MEF2C, **(C)** anti-ND4 and **(D)** anti-DRP1 after intrathecal injection of *Mef2c*-KD virus to mice. Scale bars (white): 5 µm. **(E)** Densitometry analysis showed the decrease of MEF2C and ND4 and increase of DRP1 immunoreactivity levels in GFP^+^ cells in *Mef2c*-KD mice. A total of 62 cells count, 8 cells/mouse, *N* = 4 (shControl, shMef2c). *\*P* < 0.05, *\*\*P* < 0.01; Student’s t test (*n* = 32/group, *df* = 62). **(F)** Lumbar spinal cord tissue sections were stained with cresyl violet. Scale bars: 20µm. Right panel shows quantitative analysis for motor neurons size in ventral horn. A total of 88 cells count, 11 cells/mouse, *N* = 4 (shControl, shMef2c). *\*\*\*P* < 0.001; Student’s t test (*n* = 44/group, *df* = 86). Error bars indicate means ± s.e.m.

**Supplementary Fig. 12.**
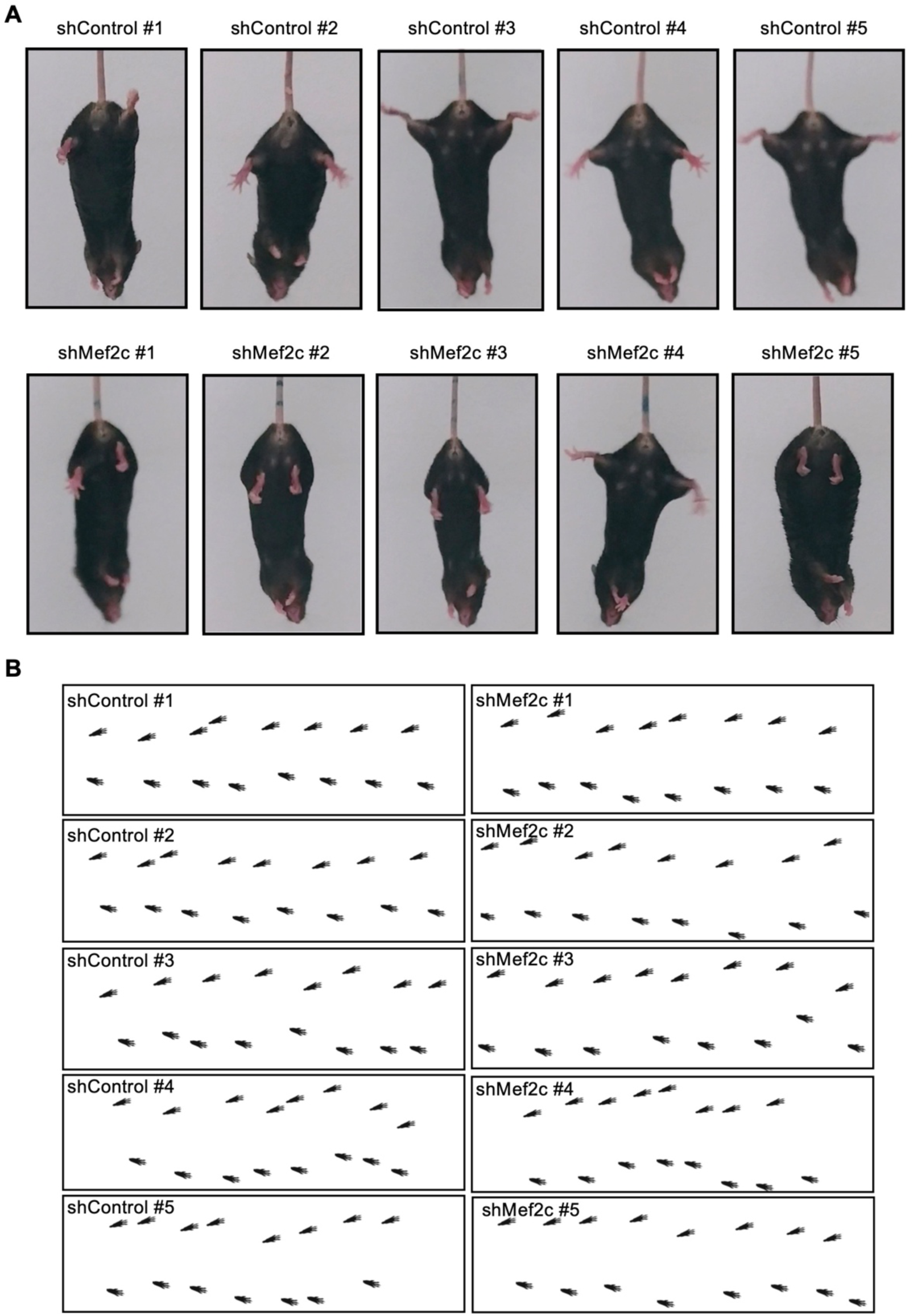
Tail suspension and gait analysis of cortical layer V *Mef2c*-KD injected mice. **(A)** Still images of tail suspension of shMef2c mice which showed abnormal hindlimb extension reflex compared to control mice. **(B)** The computer-assisted footprint in accelerated wheel running test.

**Supplementary Fig. 13.**
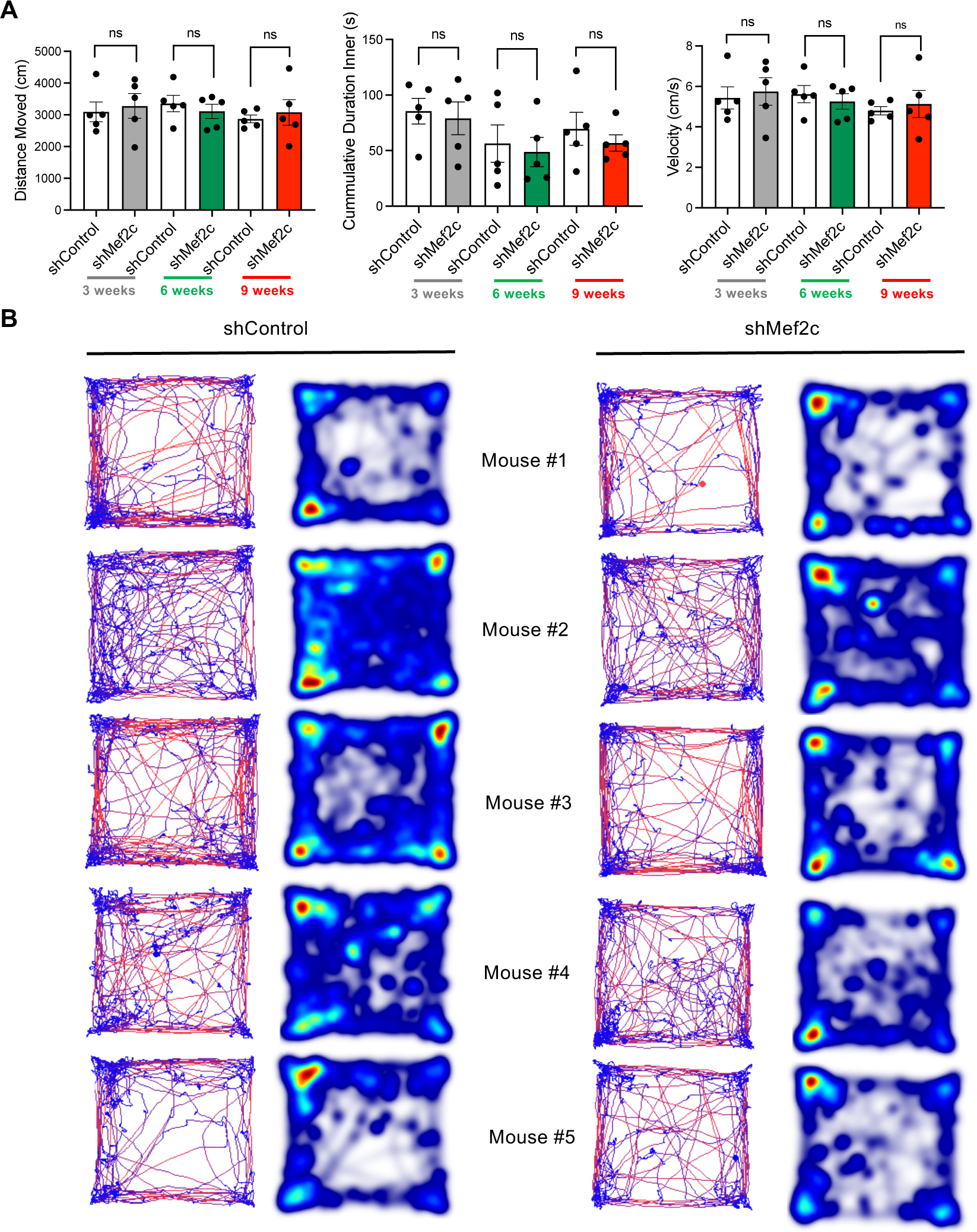
Open field test analysis results of cortical layer V *Mef2c*-KD injected mice. **(A)** Comparing distance moved, cumulative duration and velocity of shControl and shMef2c mice in open-field test 3-, 6- and 9-weeks post-injection. *N* = 5 (shControl, shMef2c). Repeated measures ANOVA test (*df* = 22). Error bars indicate means ± s.e.m. **(B)** Variable being measured in each case. Left: track visualization used to obtain the total distance traveled of mouse (red shows the higher velocity). Right: heatmap visualization.

**Supplementary Fig. 14.**
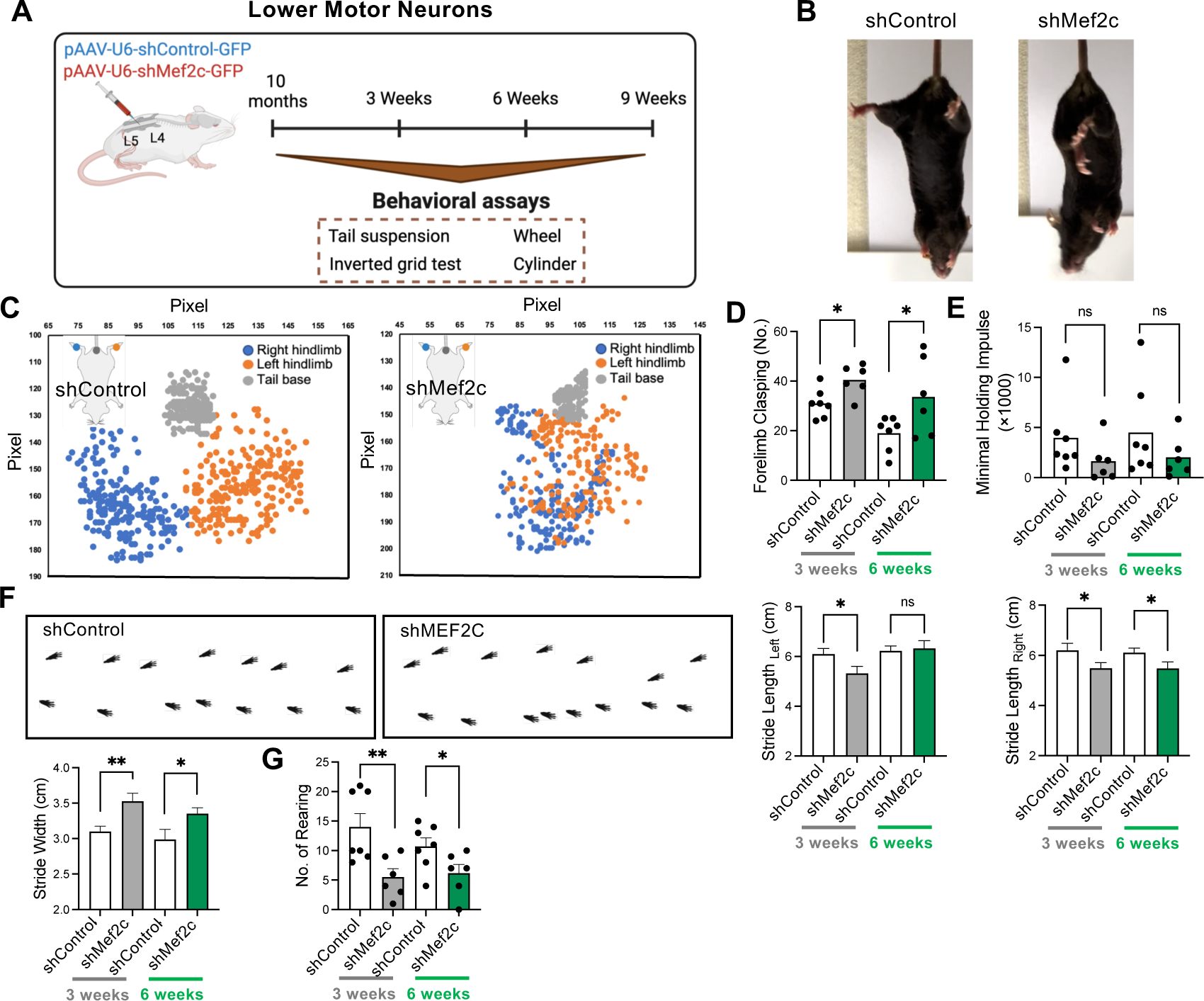
Longitudinal behavioral study for *Mef2c*-KD in the lumbar spinal cord of mice. **(A)** A scheme illustrating intrathecal injection of *Mef2c*-KD virus at L4-L5 intervertebral space and performing behavioral study 3-, 6- and 9-weeks after injection. **(B)** Still images of representative sh*Mef2c* mouse exhibited hindlimb clasping posture in tail-suspension test. **(C)** Aggregated coordinate plots in the first 10 seconds of tail suspension of shControl and shMef2c representative mice at 3-weeks post-injection. **(D)** Forelimbs clasping frequency on tail suspension test at 3- and 6-weeks post-injection. *N* = 7 shControl and 6 shMef2c mice. **(E)** Minimal holding impulse in inverted grid test decreased for sh*Mef2c* mice. Repeated measures ANOVA test (*df* = 22). **(F)** Computer-assisted footprint in wheel running test for a representative mouse in each group. Right: gait analysis showed wider stride width and shorter stride length in sh*Mef2c* mice. Repeated measures ANOVA test (*n* = 30/group, *df* = 116). **(G)** Number of rearing in cylinder test for shControl and shMef2c mice. *N* = 7 (shControl), and 6 (shMef2c). *\*P* < 0.05, *\*\*P* < 0.01; Repeated measures ANOVA test (*df* = 22). Error bars indicate means ± s.e.m.

**Supplementary Fig. 15.**
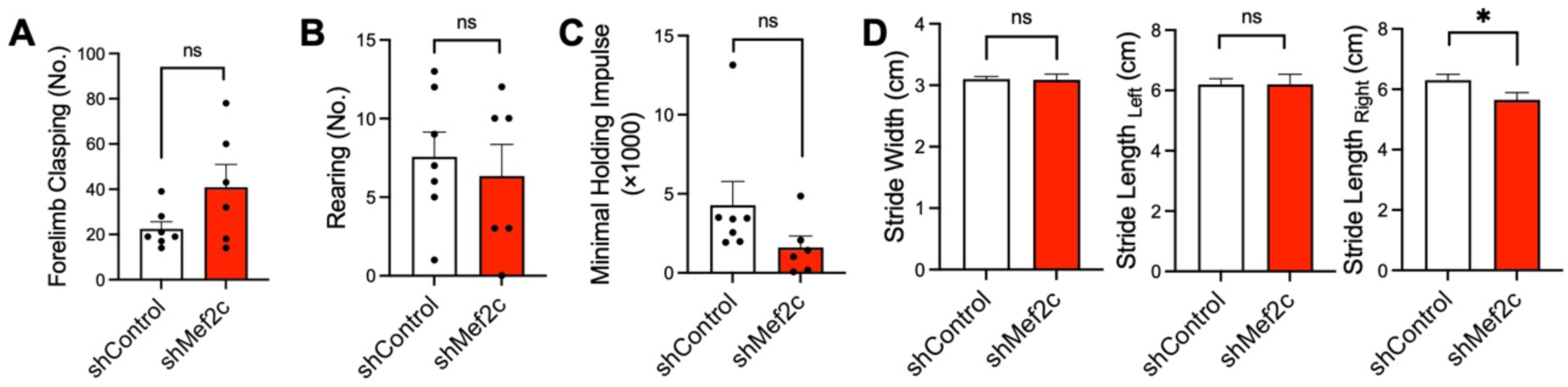
Behavioral tests at 9-weeks after intrathecal *Mef2c*-KD injection in mice. **(A)** Number of forelimbs clasping in tail suspension test. **(B)** Number of rearing in cylinder test. **(C)** Minimal holding impulse in inverted grid test. Student’s t test (*df* = 11). **(D)** Gait analysis in accelerated wheel test. *N* = 7 (shControl), and 6 (shMef2c). *\*P* < 0.05; Student’s t test (*n* = 40/group, *df* = 78). Error bars indicate means ± s.e.m.

**Supplementary Fig. 16.**
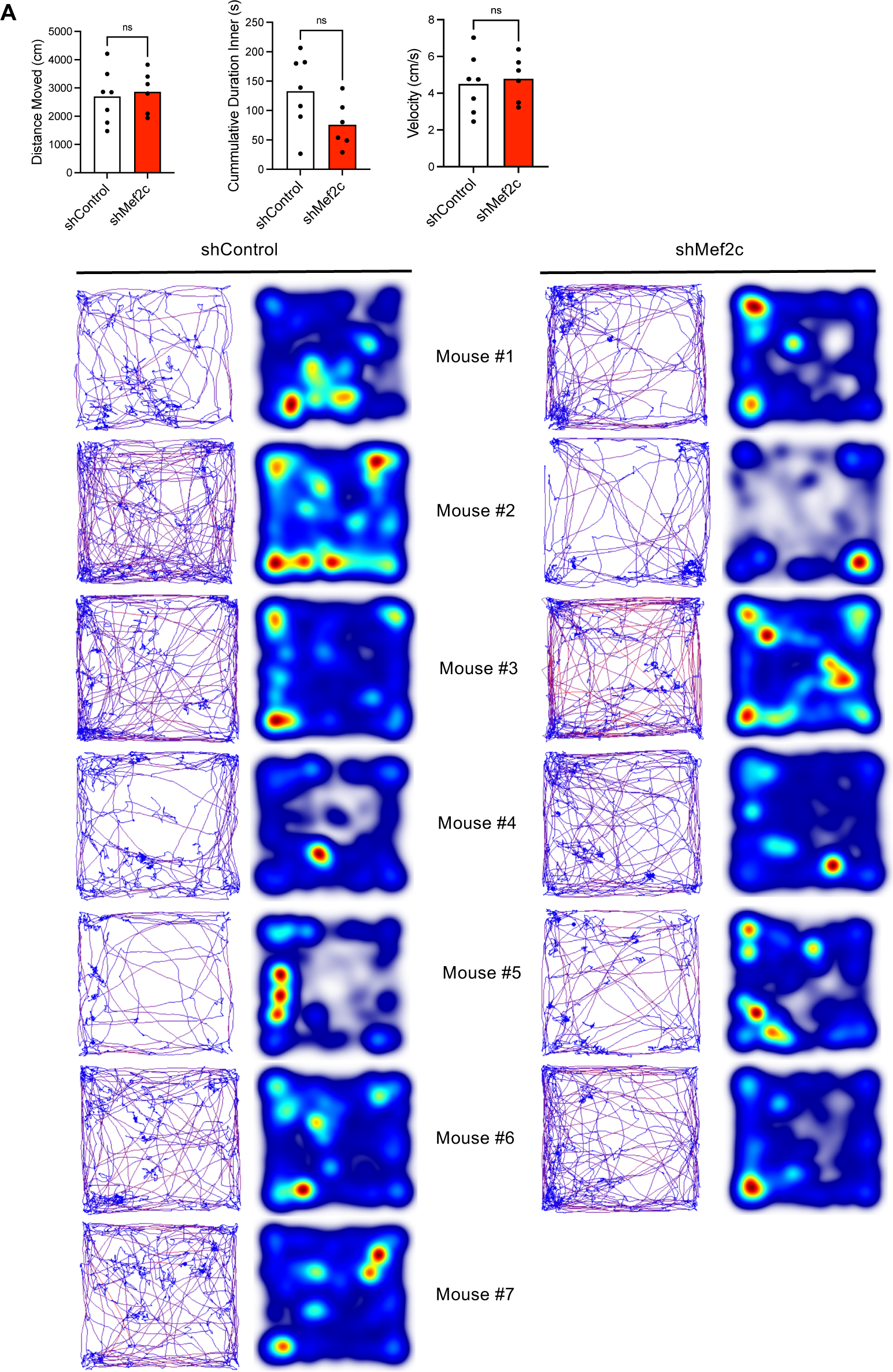
Open field test analysis results of intrathecal *Mef2c*-KD injected mice. **(A)** Comparing distance moved, cumulative duration and velocity of shControl and shMef2c mice in open-field test 3-weeks post-injection. *N* = 7 (shControl), and 6 (shMef2c). Student’s t test (*df* = 11). Error bars indicate means ± s.e.m. **(B)** Variable being measured in each case. Left: track visualization used to obtain the total distance traveled of mouse (red shows the higher velocity). Right: heatmap visualization.

**Supplementary Fig. 17.**
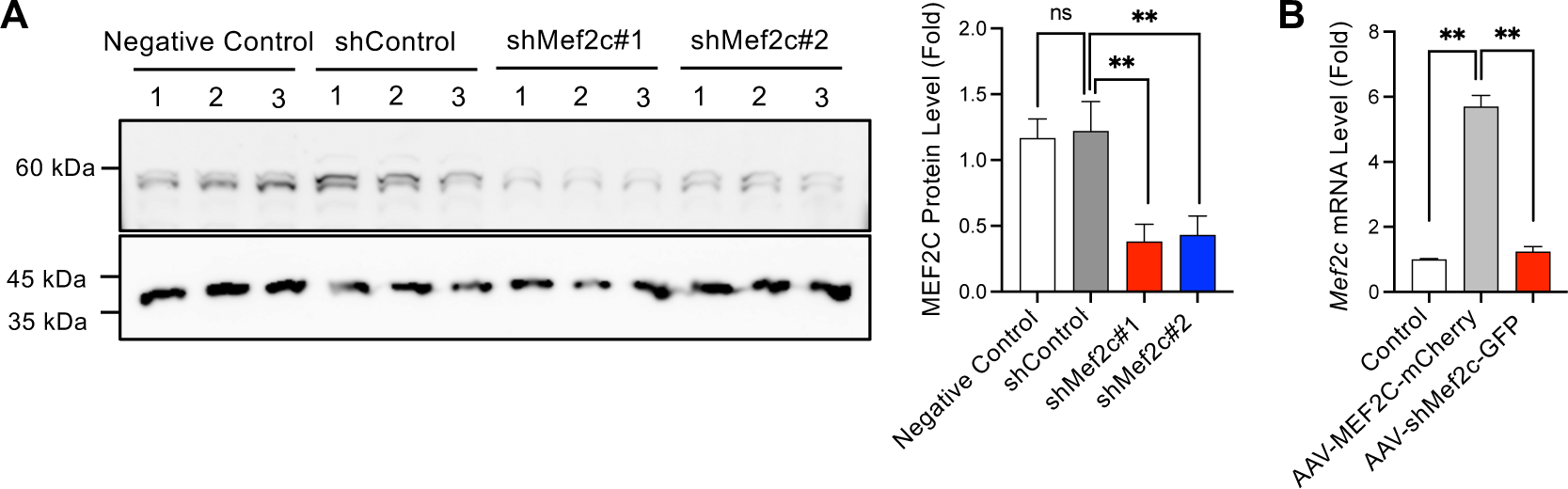
Verify the efficiency of *Mef2c*-KD and *MEF2C*-O/E viruses in NSC-34 cell line. **(A)** NSC-34 cells were infected with shMef2c#1 (used in this study), shMef2c#2 or shControl for 48 hrs and protein was extracted with RIPA. Right: western blot quantitation of MEF2C level. *\*\*P* < 0.01; One-way ANOVA test (*n* = 3/group, *df* = 8). **(B)** qPCR results showed *Mef2c* mRNA level in NSC-34 cells infected by AAV-MEF2C-mCherry (overexpression) and AAV-shMef2c-GFP (knockdown) viruses. *\*\*P* < 0.01; Student’s t test. One-way ANOVA test (*n* = 2/group, *df* = 3). Error bars indicate means ± s.e.m.

**Supplementary Table 1.** The 27 SNPs predicated as the candidate risk-SNPs and -genes for ALS.

| **Gene** | **SNP** | **Location** | **GWAS *p*-value** |
| --- | --- | --- | --- |
| CLCN3, NEK1,<br>C4orf27 | rs11722212 | chr4:170534428 | 6.79E-05 |
| MEF2C | rs700587 | chr5:88110084 | 2.82E-04 |
| MEF2C | rs304153 | chr5:88121188 | 3.65E-04 |
| MEF2C | rs304152 | chr5:88124123 | 3.75E-04 |
| MEF2C | rs304151 | chr5:88125853 | 3.19E-04 |
| NEK1 | rs7687276 | chr4:170319471 | 3.55E-05 |
| NEK1 | rs6827421 | chr4:170326359 | 1.82E-05 |
| NEK1 | rs6831487 | chr4:170331972 | 1.46E-05 |
| MSRA | rs11989640 | chr8:10256054 | 3.59E-04 |
| TENM2 | rs4242226 | chr5:167181730 | 2.90E-05 |
| FOXP1 | rs9827299 | chr3:71353372 | 4.84E-04 |
| FOXP1 | rs76753627 | chr3:71356015 | 4.91E-04 |
| SDCCAG8 | rs58560561 | chr1:243537729 | 2.54E-04 |
| RBFOX1 | rs11077038 | chr16:6460736 | 3.85E-04 |
| ZNHIT3 | rs4796224 | chr17:34842521 | 3.66E-05 |
| NFIB | rs7021539 | chr9:14282792 | 4.46E-04 |
| PTPRD | rs10756021 | chr9:10292307 | 2.32E-04 |
| ZNF184 | rs7744110 | chr6:27421304 | 6.49E-05 |
| ZNF184 | rs2235252 | chr6:27422493 | 6.66E-05 |
| ZNF184 | rs2235254 | chr6:27422794 | 6.55E-05 |
| ZNF184 | rs12199110 | chr6:27423568 | 6.37E-05 |
| ZNF184 | rs2092121 | chr6:27424184 | 6.66E-05 |
| ZNF184 | rs2092122 | chr6:27424311 | 6.66E-05 |
| ZNF184 | rs764460 | chr6:27424374 | 6.43E-05 |
| ZNF184 | rs764461 | chr6:27424443 | 6.60E-05 |
| ZNF184 | rs10484398 | chr6:27424960 | 6.40E-05 |
| ZNF184 | rs6456789 | chr6:27426498 | 6.61E-05 |

**Supplementary Table 2.** Information of human tissues from normal subjects and ALS patients.

| **Number** | **Case (Normal or ALS)** | **Familial or Sporadic ALS** | **Sex** | **Age** | **PMI** |
| --- | --- | --- | --- | --- | --- |
| 1 | Normal |  | M | 70 | <24 |
| 2 | Normal |  | M | 84 | <24 |
| 3 | Normal |  | M | 60 | <24 |
| 4 | Normal |  | M | 69 | <24 |
| 5 | Normal |  | M | 60 | <24 |
| 1 | ALS | Sporadic | M | 84 | 27.5 |
| 2 | ALS | Sporadic | M | 45 | 51.6 |
| 3 | ALS | Sporadic | M | 60 | 43.6 |
| 4 | ALS | Sporadic | M | 78 | 25.7 |
| 5 | ALS | Sporadic | M | 62 | 19.4 |

**Supplementary Table 3.**
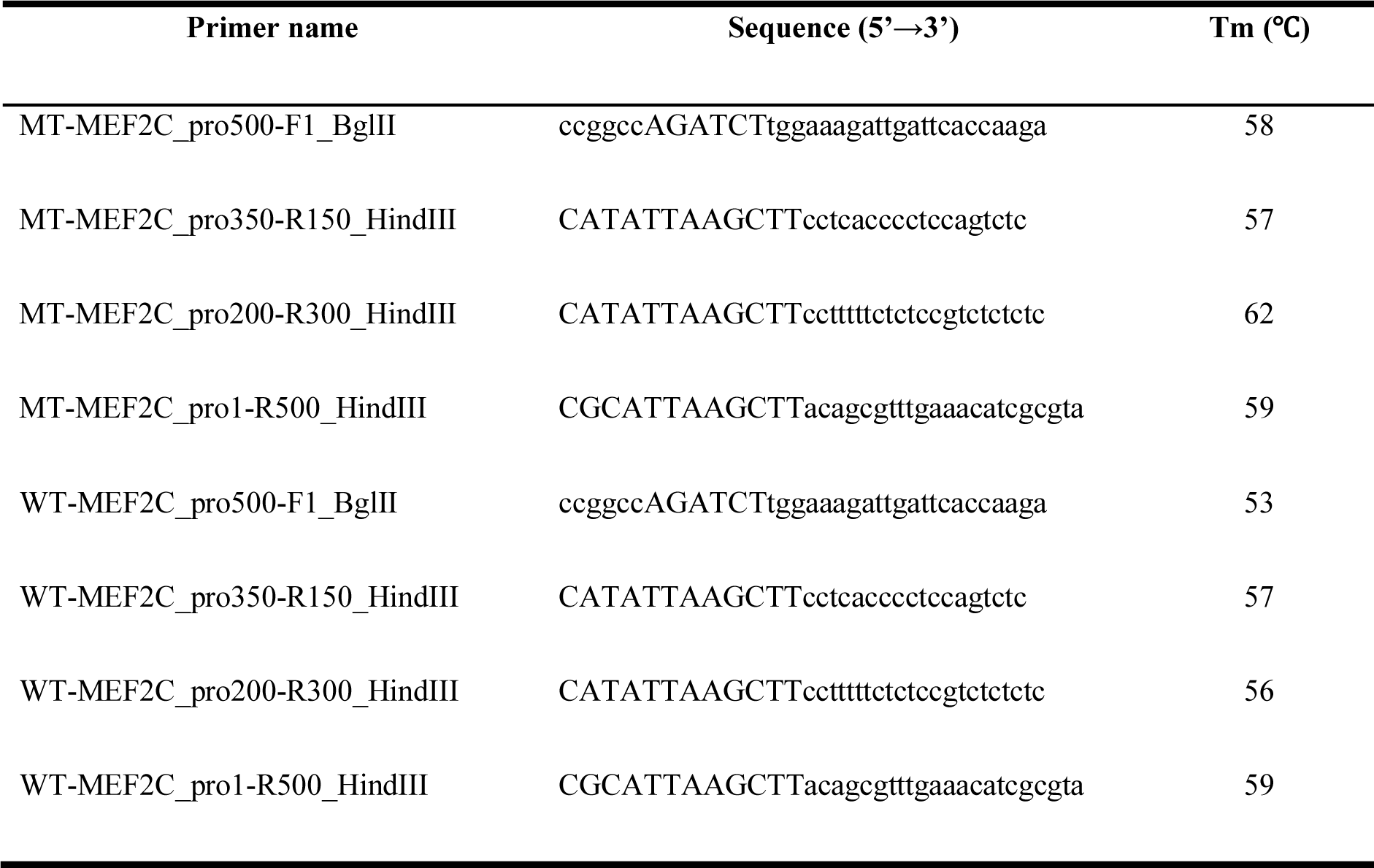
PCR primers that were used for pGL4.14-duplex-MEF2C promoter cloning.

**Supplementary Table 4.**
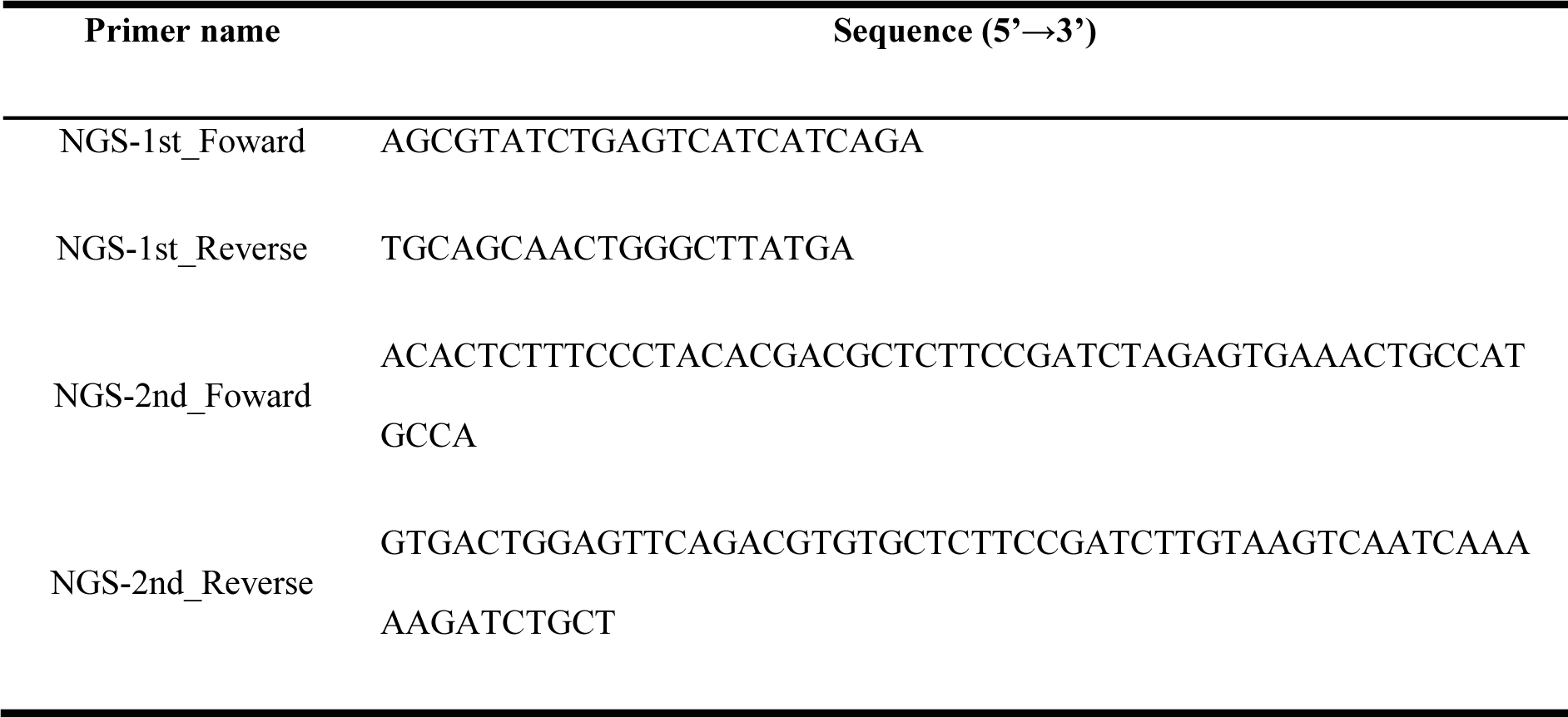
PCR primers for NGS library generation.

**Supplementary Table 5.**
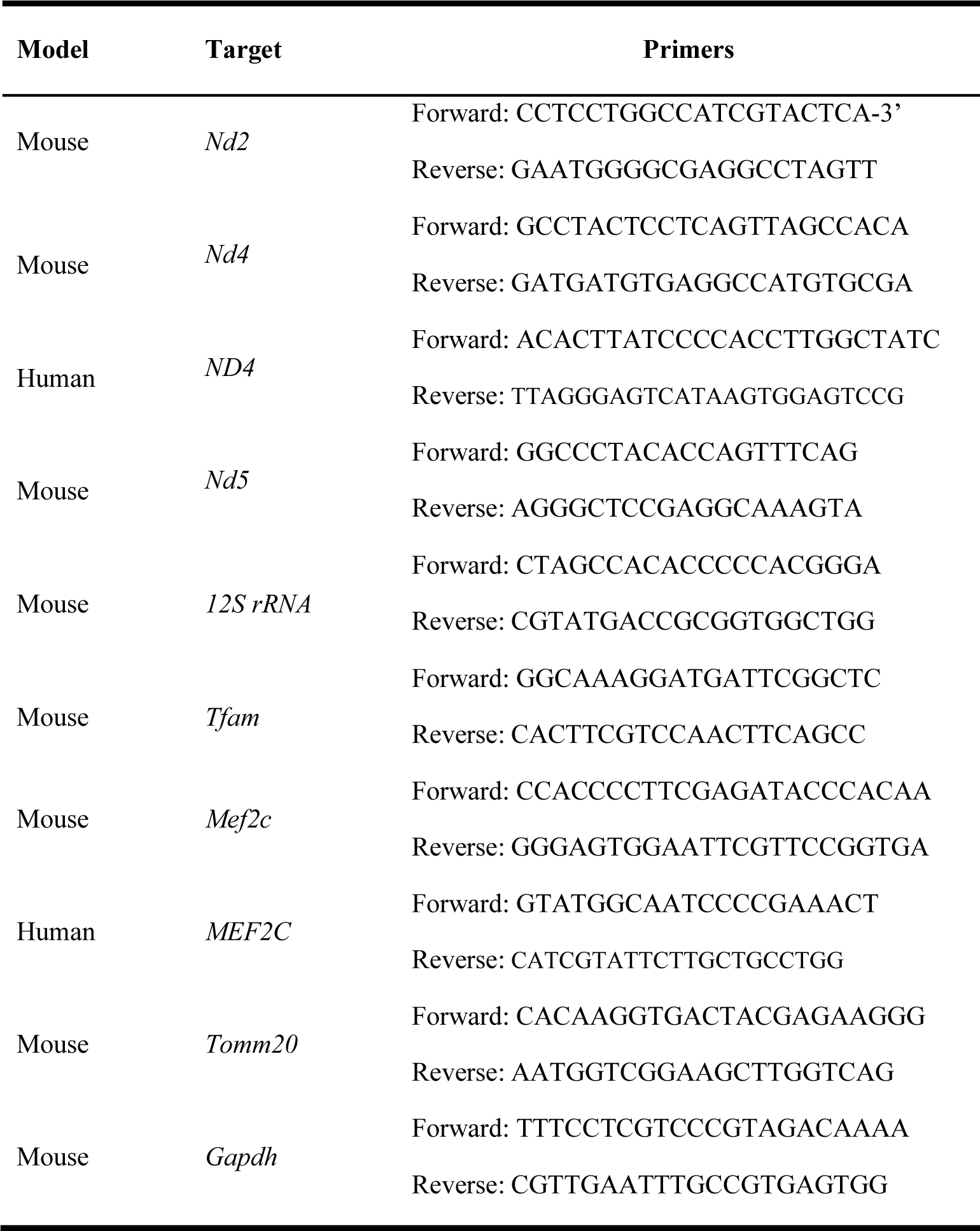
qPCR primers that were used for determining the relative levels of gene expression.

**Supplementary Table 6.**
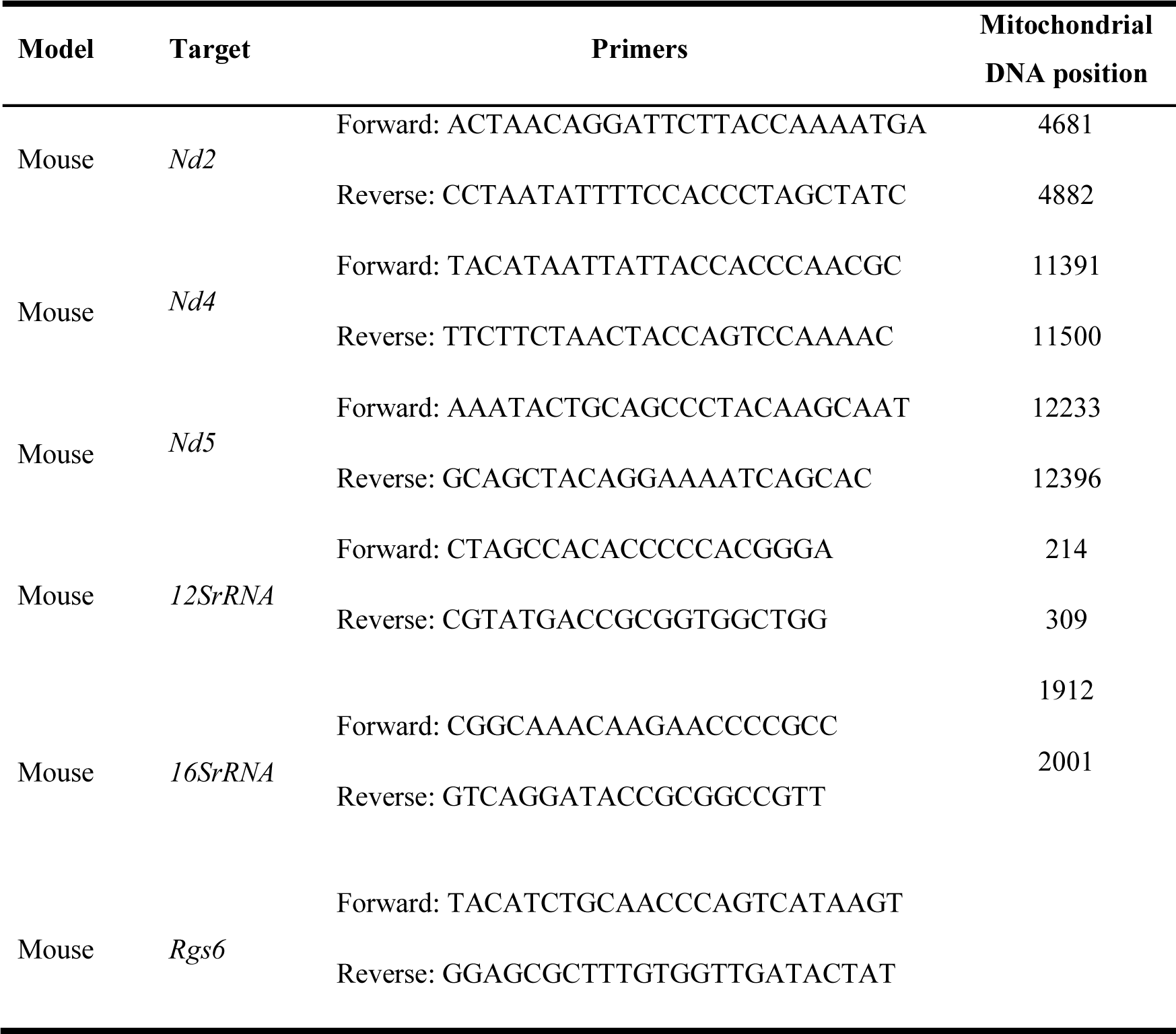
qPCR primers that were used for MEF2C ChIP-qPCR.

**Supplementary Table 7.**
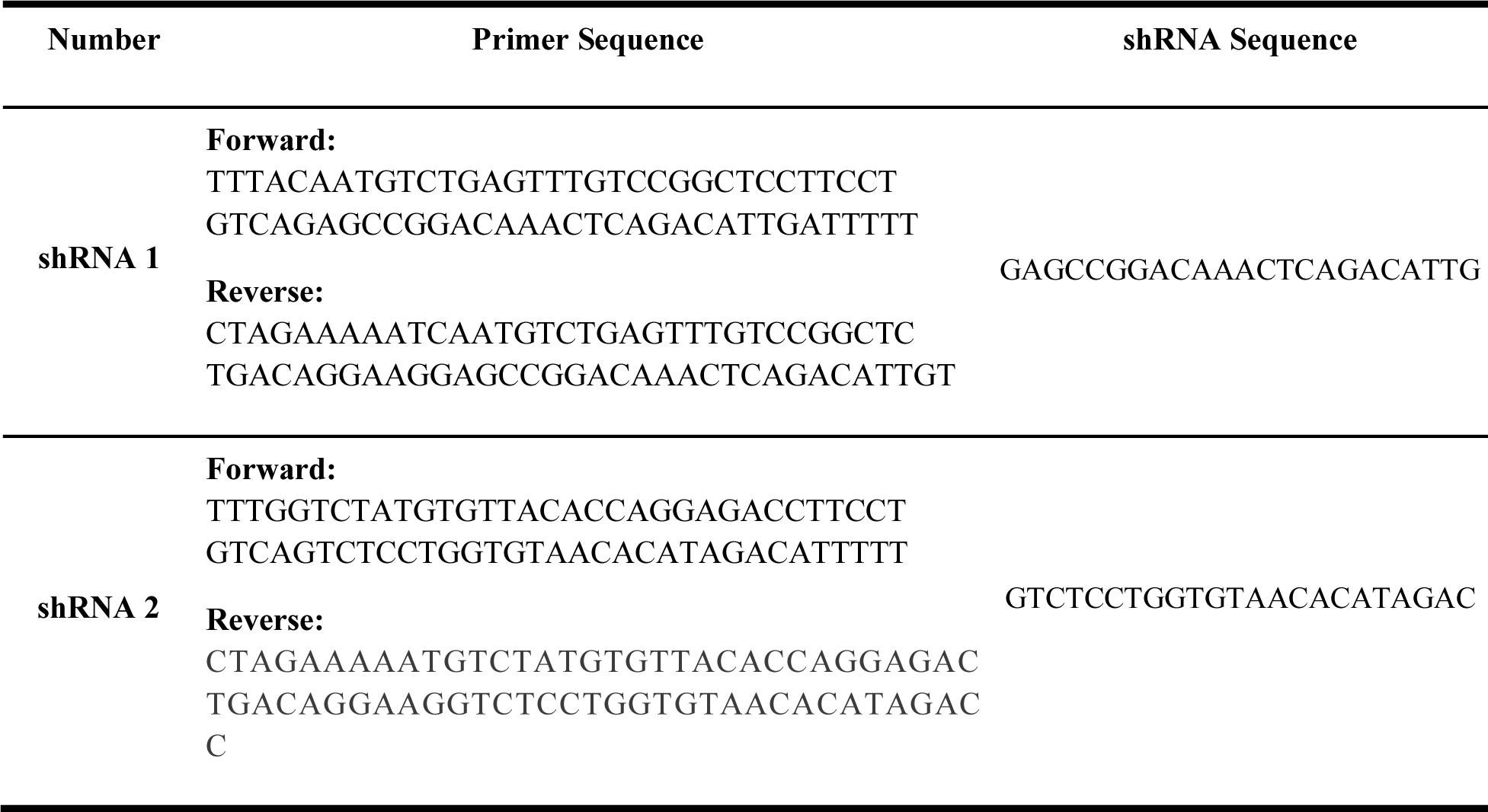
Primer design sequences for shRNA-Mef2c plasmids.

## References

1. Haukedal H, Freude K. Implications of Microglia in Amyotrophic Lateral Sclerosis and Frontotemporal Dementia. Journal of Molecular Biology. 2019;431(9):1818–29.

2. Yousefian-Jazi A, Seol Y, Kim J, Ryu HL, Lee J, Ryu H. Pathogenic Genome Signatures That Damage Motor Neurons in Amyotrophic Lateral Sclerosis. Cells. 2020;9(12):2687.

3. Belzil VV, Bauer PO, Prudencio M, Gendron TF, Stetler CT, Yan IK, et al. Reduced C9orf72 gene expression in c9FTD/ALS is caused by histone trimethylation, an epigenetic event detectable in blood. Acta Neuropathol. 2013;126(6):895–905.

4. Belzil VV, Bauer PO, Gendron TF, Murray ME, Dickson D, Petrucelli L. Characterization of DNA hypermethylation in the cerebellum of c9FTD/ALS patients. Brain Res. 2014;1584:15–21.

5. Xi Z, Rainero I, Rubino E, Pinessi L, Bruni AC, Maletta RG, et al. Hypermethylation of the CpG-island near the C9orf72 G₄C₂-repeat expansion in FTLD patients. Hum Mol Genet. 2014;23(21):5630–7.

6. Xi Z, Zhang M, Bruni AC, Maletta RG, Colao R, Fratta P, et al. The C9orf72 repeat expansion itself is methylated in ALS and FTLD patients. Acta Neuropathol. 2015;129(5):715–27.

7. Bennett SA, Tanaz R, Cobos SN, Torrente MP. Epigenetics in amyotrophic lateral sclerosis: a role for histone post-translational modifications in neurodegenerative disease. Translational Research. 2019;204:19–30.

8. Choi SH, Yousefian-Jazi A, Hyeon SJ, Nguyen PTT, Chu J, Kim S, et al. Modulation of histone H3K4 dimethylation by spermidine ameliorates motor neuron survival and neuropathology in a mouse model of ALS. J Biomed Sci. 2022;29(1):106.

9. Chen Z, Rasheed M, Deng Y. The epigenetic mechanisms involved in mitochondrial dysfunction: Implication for Parkinson’s disease. Brain Pathol. 2022;32(3):e13012.

10. van Rheenen W, Shatunov A, Dekker AM, McLaughlin RL, Diekstra FP, Pulit SL, et al. Genome-wide association analyses identify new risk variants and the genetic architecture of amyotrophic lateral sclerosis. Nature Genetics. 2016;48(9):1043–8.

11. Fogh I, Ratti A, Gellera C, Lin K, Tiloca C, Moskvina V, et al. A genome-wide association meta-analysis identifies a novel locus at 17q11.2 associated with sporadic amyotrophic lateral sclerosis. Hum Mol Genet. 2014;23(8):2220–31.

12. Nicolas A, Kenna KP, Renton AE, Ticozzi N, Faghri F, Chia R, et al. Genome-wide Analyses Identify KIF5A as a Novel ALS Gene. Neuron. 2018;97(6):1268–83.e6.

13. Tam V, Patel N, Turcotte M, Bossé Y, Paré G, Meyre D. Benefits and limitations of genome-wide association studies. Nature Reviews Genetics. 2019;20(8):467–84.

14. Lee T, Sung MK, Lee S, Yang W, Oh J, Kim JY, et al. Convolutional neural network model to predict causal risk factors that share complex regulatory features. Nucleic Acids Research. 2019;47(22):e146-e.

15. Yousefian-Jazi A, Sung MK, Lee T, Hong Y-H, Choi JK, Choi J. Functional fine-mapping of noncoding risk variants in amyotrophic lateral sclerosis utilizing convolutional neural network. Scientific Reports. 2020;10(1):12872.

16. Ripke S, Neale BM, Corvin A, Walters JTR, Farh K-H, Holmans PA, et al. Biological insights from 108 schizophrenia-associated genetic loci. Nature. 2014;511(7510):421-7.

17. Cao Q, Anyansi C, Hu X, Xu L, Xiong L, Tang W, et al. Reconstruction of enhancer–target networks in 935 samples of human primary cells, tissues and cell lines. Nature Genetics. 2017;49(10):1428–36.

18. McLaughlin RL, Schijven D, van Rheenen W, van Eijk KR, O’Brien M, Kahn RS, et al. Genetic correlation between amyotrophic lateral sclerosis and schizophrenia. Nature Communications. 2017;8(1):14774.

19. Scekic-Zahirovic J, Sanjuan-Ruiz I, Kan V, Megat S, De Rossi P, Dieterlé S, et al. Cytoplasmic FUS triggers early behavioral alterations linked to cortical neuronal hyperactivity and inhibitory synaptic defects. Nature Communications. 2021;12(1):3028.

20. Stranger BE, Brigham LE, Hasz R, Hunter M, Johns C, Johnson M, et al. Enhancing GTEx by bridging the gaps between genotype, gene expression, and disease. Nature Genetics. 2017;49(12):1664–70.

21. Yang D, Jang I, Choi J, Kim M-S, Lee AJ, Kim H, et al. 3DIV: A 3D-genome Interaction Viewer and database. Nucleic Acids Research. 2017;46(D1):D52–D7.

22. Fahey L, Ali D, Donohoe G, Ó Broin P, Morris DW. Genes positively regulated by Mef2c in cortical neurons are enriched for common genetic variation associated with IQ and educational attainment. Human Molecular Genetics. 2023;32(22):3194–203.

23. Lee J, Kim Y, Liu T, Hwang YJ, Hyeon SJ, Im H, et al. SIRT3 deregulation is linked to mitochondrial dysfunction in Alzheimer’s disease. Aging Cell. 2018;17(1):e12679.

24. Hu D, Qi X. Quantifying Drp1-Mediated Mitochondrial Fission by Immunostaining in Fixed Cells. Methods Mol Biol. 2020;2159:197–204.

25. Liu J, Yang M, Su M, Liu B, Zhou K, Sun C, et al. FOXG1 sequentially orchestrates subtype specification of postmitotic cortical projection neurons. Science Advances. 2022;8(21):eabh3568.

26. Hirai T, Enomoto M, Kaburagi H, Sotome S, Yoshida-Tanaka K, Ukegawa M, et al. Intrathecal AAV Serotype 9-mediated Delivery of shRNA Against TRPV1 Attenuates Thermal Hyperalgesia in a Mouse Model of Peripheral Nerve Injury. Molecular Therapy. 2014;22(2):409–19.

27. Rivera Chloe M, Ren B. Mapping Human Epigenomes. Cell. 2013;155(1):39–55.

28. Lange PS, Chavez JC, Pinto JT, Coppola G, Sun C-W, Townes TM, et al. ATF4 is an oxidative stress–inducible, prodeath transcription factor in neurons in vitro and in vivo. Journal of Experimental Medicine. 2008;205(5):1227–42.

29. Palstra R-J. Transcription factor binding at enhancers: shaping a genomic regulatory landscape in flux. Front Genet. 2012;3.

30. Janson CG, Chen Y, Li Y, Leifer D. Functional regulatory regions of human transcription factor MEF2C. Brain Res Mol Brain Res. 2001;97(1):70–82.

31. Zhao M, New L, Kravchenko VV, Kato Y, Gram H, di Padova F, et al. Regulation of the MEF2 family of transcription factors by p38. Mol Cell Biol. 1999;19(1):21–30.

32. Satyaraj E, Storb U. Mef2 Proteins, Required for Muscle Differentiation, Bind an Essential Site in the Ig λ Enhancer1. The Journal of Immunology. 1998;161(9):4795–802.

33. Beal MF. Mitochondria take center stage in aging and neurodegeneration. Ann Neurol. 2005;58(4):495–505.

34. Kang I, Chu CT, Kaufman BA. The mitochondrial transcription factor TFAM in neurodegeneration: emerging evidence and mechanisms. FEBS letters. 2018;592(5):793–811.

35. Needs HI, Protasoni M, Henley JM, Prudent J, Collinson I, Pereira GC. Interplay between Mitochondrial Protein Import and Respiratory Complexes Assembly in Neuronal Health and Degeneration. Life (Basel). 2021;11(5).

36. Vayssiere JL, Petit PX, Risler Y, Mignotte B. Commitment to apoptosis is associated with changes in mitochondrial biogenesis and activity in cell lines conditionally immortalized with simian virus 40. Proc Natl Acad Sci U S A. 1994;91(24):11752–6.

37. Muyderman H, Chen T. Mitochondrial dysfunction in amyotrophic lateral sclerosis - a valid pharmacological target? Br J Pharmacol. 2014;171(8):2191–205.

38. Mitchell AC, Javidfar B, Pothula V, Ibi D, Shen EY, Peter CJ, et al. MEF2C transcription factor is associated with the genetic and epigenetic risk architecture of schizophrenia and improves cognition in mice. Molecular Psychiatry. 2018;23(1):123–32.

39. Udeochu JC, Amin S, Huang Y, Fan L, Torres ERS, Carling GK, et al. Tau activation of microglial cGAS–IFN reduces MEF2C-mediated cognitive resilience. Nature Neuroscience. 2023;26(5):737–50.

40. Li W-K, Zhang S-Q, Peng W-L, Shi Y-H, Yuan B, Yuan Y-T, et al. Whole-brain in vivo base editing reverses behavioral changes in Mef2c-mutant mice. Nature Neuroscience. 2023.

41. Claringbould A, Zaugg JB. Enhancers in disease: molecular basis and emerging treatment strategies. Trends in Molecular Medicine. 2021;27(11):1060–73.

42. Preisig DF, Kulic L, Krüger M, Wirth F, McAfoose J, Späni C, et al. High-speed video gait analysis reveals early and characteristic locomotor phenotypes in mouse models of neurodegenerative movement disorders. Behavioural Brain Research. 2016;311:340–53.

43. Yoon Y, Park H, Kim S, Nguyen PT, Hyeon SJ, Chung S, et al. Genetic Ablation of EWS RNA Binding Protein 1 (EWSR1) Leads to Neuroanatomical Changes and Motor Dysfunction in Mice. Exp Neurobiol. 2018;27(2):103–11.

44. Li X, Chen C, Zhan X, Li B, Zhang Z, Li S, et al. R13 preserves motor performance in SOD1^G93A^ mice by improving mitochondrial function. Theranostics. 2021;11(15):7294–307.

45. Trabjerg MS, Andersen DC, Huntjens P, Oklinski KE, Bolther L, Hald JL, et al. Downregulating carnitine palmitoyl transferase 1 affects disease progression in the SOD1 G93A mouse model of ALS. Communications Biology. 2021;4(1):509.

46. Wainger BJ, Kiskinis E, Mellin C, Wiskow O, Han SS, Sandoe J, et al. Intrinsic membrane hyperexcitability of amyotrophic lateral sclerosis patient-derived motor neurons. Cell Rep. 2014;7(1):1–11.

47. Kuo JJ, Schonewille M, Siddique T, Schults ANA, Fu R, Bär PR, et al. Hyperexcitability of Cultured Spinal Motoneurons From Presymptomatic ALS Mice. Journal of Neurophysiology. 2004;91(1):571–5.

48. Heinemann U, Lux HD. Undershoots following stimulus-induced rises of extracellular potassium concentration in cerebral cortex of cat. Brain Res. 1975;93(1):63–76.

49. Nimchinsky EA, Yasuda R, Oertner TG, Svoboda K. The number of glutamate receptors opened by synaptic stimulation in single hippocampal spines. J Neurosci. 2004;24(8):2054–64.

50. Abramov AY, Duchen MR. Impaired mitochondrial bioenergetics determines glutamate-induced delayed calcium deregulation in neurons. Biochim Biophys Acta. 2010;1800(3):297–304.

51. Kann O, Kovács R. Mitochondria and neuronal activity. American Journal of Physiology-Cell Physiology. 2007;292(2):C641–C57.

52. Schreiber J, Singh R, Bilmes J, Noble WS. A pitfall for machine learning methods aiming to predict across cell types. bioRxiv. 2019:512434.

53. Lee J, Lim K, Kim A, Mok YG, Chung E, Cho S-I, et al. Prime editing with genuine Cas9 nickases minimizes unwanted indels. Nature Communications. 2023;14(1):1786.

